# ErbB Signalling is a Potential Therapeutic Target for Vascular Lesions with Fibrous Component

**DOI:** 10.1101/2022.09.23.509204

**Authors:** H. Ilmonen, S. Jauhiainen, P. Vuola, H. Rasinkangas, H.H. Pulkkinen, S. Keränen, M. Kiema, J.J. Liikkanen, N. Laham-Karam, S. Laidinen, E. Aavik, K. Lappalainen, J. Lohi, J. Aronniemi, T. Örd, M.U. Kaikkonen, P. Salminen, E. Tukiainen, S. Ylä-Herttuala, J.P. Laakkonen

## Abstract

**Background:** Sporadic venous malformation (VM) and angiomatosis of soft tissue (AST) are benign, congenital vascular anomalies affecting venous vasculature. Depending on the size and location of the lesion, symptoms vary from motility disturbances to pain and disfigurement. Due to high recurrence of the lesions more effective therapies are needed.

**Methods:** As targeting stromal cells has been an emerging concept in anti-angiogenic therapies, here, by using VM/AST patient samples, RNA-sequencing, cell culture techniques and a xenograft mouse model, we investigated the crosstalk of endothelial cells (EC) and fibroblasts and its effect on vascular lesion growth.

**Results:** We report, for the first time, expression and secretion of transforming growth factor A (TGFA) in ECs or intervascular stromal cells in AST and VM lesions. TGFA induced secretion of VEGF-A paracrinally, and regulated EC proliferation. Oncogenic PIK3CA variant in p.H1047R, a common somatic mutation found in these lesions, increased TGFA expression, enrichment of hallmark hypoxia, and in a mouse xenograft model, lesion size and vascularization. Treatment with afatinib, a pan-ErbB tyrosine-kinase inhibitor, decreased vascularization and lesion size in mouse xenograft model with ECs expressing oncogenic PIK3CA p.H1047R variant and fibroblasts.

**Conclusions:** Based on the data, we suggest that targeting of both intervascular stromal cells and ECs is a potential treatment strategy for vascular lesions having a fibrous component.

**Funding:** Academy of Finland, Ella and Georg Ehnrooth foundation, the ERC grants, Sigrid Jusélius Foundation, Finnish Foundation for Cardiovascular Research, Jane and Aatos Erkko Foundation, and Department of Musculosceletal and Plastic Surgery, Helsinki University Hospital.

**GRAPHICAL ABSTRACT:** Graphical abstract. Proposed model for the paracrine signaling of TGFA/VEGF-A in vascular lesion
Schematic illustration showing the general structure of venous malformation or angiomatosis of soft tissue. Pathological vasculature in the lesion (dark blue) is surrounded by disorganized extracellular matrix (ECM) and intervascular stromal cells (SCs, orange). High magnification from the area close to vessel wall demonstrates the proposed model for crosstalk between endothelial cells (ECs) and SCs. A mutation in phosphatidylinositol-4,5-biphosphate 3-kinase catalytic subunit alpha (PIK3CA) gene **(1)** or other processes promote ECs to express high level of transforming growth factor A (TGFA) **(2)**. TGFA binds to epithelial growth factor receptor (EGFR) on the surface of adjacent SCs **(3)**. Activated EGFR-downstream signaling **(4)** promotes elevated expression of vascular endothelial growth factor (VEGF)-A in SCs and increases the expression of TGFA **(5)**. VEGF-A secreted from SCs **(6)** binds to VEGF-recetor-2 (VEGFR2) on surface of ECs **(7)** and together with TGFA activates angiogenic EC phenotype. TGFA secreted from the SCs **(8)**, can further activate EGFR and its downstream signaling.

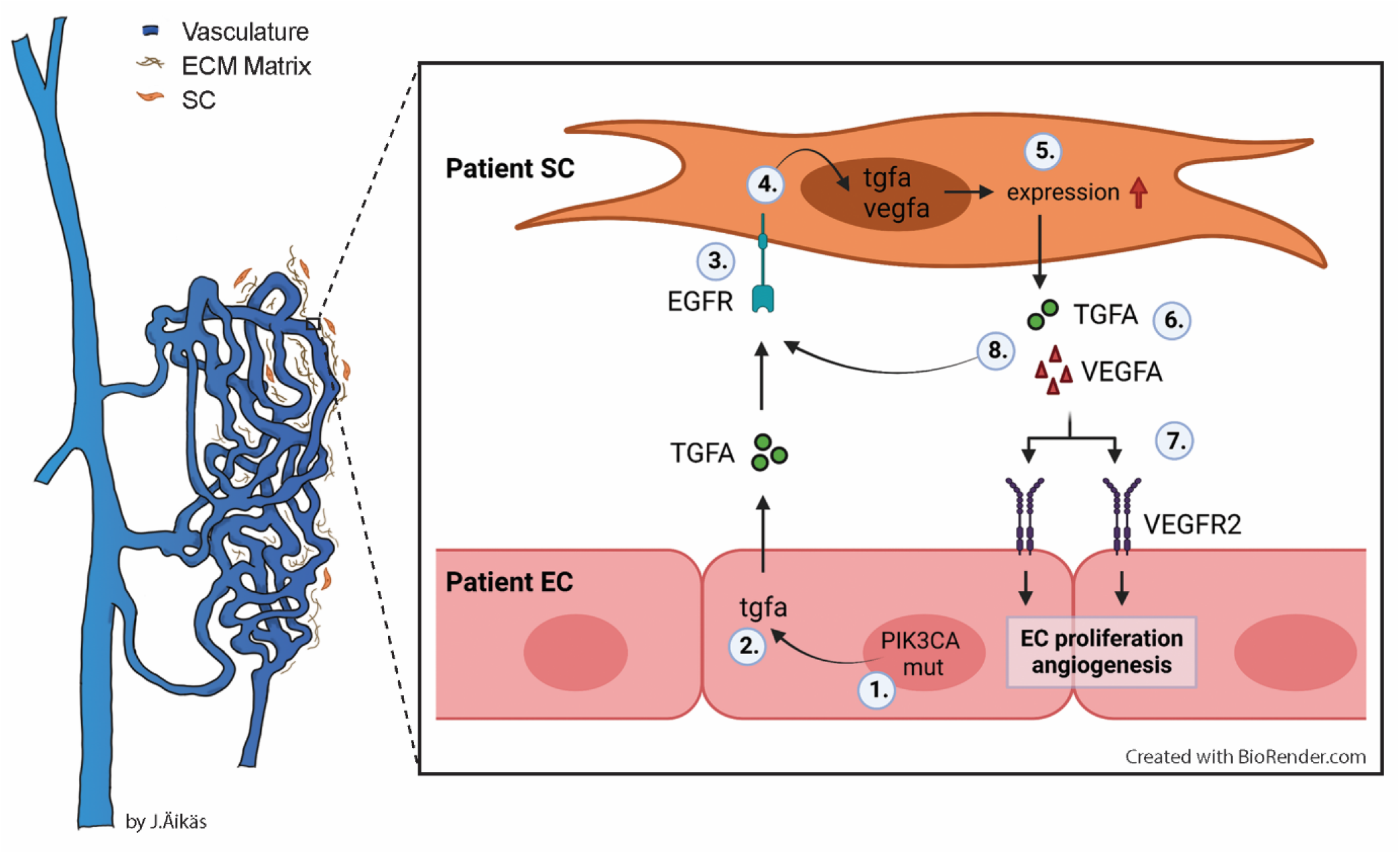

## INTRODUCTION

Sporadic venous malformation (VM) and angiomatosis of soft tissue (AST) form a heterogeneous group of vascular anomalies affecting venous vasculature (1,2). Lesions form due to a local defect in vascular development during embryogenesis and expand with time, manifesting clinically usually in late childhood or early adulthood (3). VM can locate in any tissue or internal organ and be either superficial or permeate multiple tissue planes (4), whereas AST is typically found in extremities or trunk being subcutaneous or intramuscular. In both, symptoms vary from limited aesthetic harm to motility disturbances, muscle weakness, pain, disfigurement, and life-threatening bleeding, depending on the size and location of the lesion.

Overlapping magnetic resonance imaging findings, but distinctive histological features, are usually found between VM and AST (5). In both AST and VM, venous structures form enlarged, irregular vascular channels (5,6). Fibrous connective tissue with fibroblasts is detected around the various-sized vessels in AST (5), while sclerotherapy can cause secondary fibrosis in VMs (7). Whereas VMs consists solely of venous structures, AST also has artery-like vessels, lymphatic vessels, small capillaries and mesenchymal tissue components, especially muscle-infiltrating fat (5,8) In ISSVA classification (i.e. International Society for the Study of Vascular Anomalies), AST is classified under provisionally unclassified vascular anomalies (issva.org/classification, y.2018). In sporadic VMs, somatic mutations in tyrosine protein kinase receptor TEK are found in approximately half of the patients, while somatic PIK3CA mutations are found in 20% of the VMs lacking TEK alterations (9,10). Somatic PIK3CA mutations have also been associated with AST (11). Causative mutations in these genes lead to chronic activation of AKT and dysregulation of EC migration, expression of angiogenic factors as well as alterations in composition and processing of the extracellular matrix (9,12,13)

VM or AST do not regress spontaneously. If conservative treatment is ineffective, symptomatic lesions are treated with percutaneous sclerotherapy, percutaneous cryotherapy, endovascular laser treatment or surgical resection (14–18). At present, sirolimus targeting the PI3K/AKT/mTOR pathway is tested in clinical trials for the treatment of VM (19,20) (ClinicalTrials.gov, study nro: NCT02638389). So far, most of the previous studies have been focusing on the role of ECs in VM or AST pathogenesis. As targeting of stromal cells is an emerging concept for the development of anti-angiogenic therapies (21–23), we studied here crosstalk of ECs and intervascular stromal cells in VM and AST and assessed the role of fibroblasts in PI3K-driven lesion growth in mice.

## RESULTS

### Patient Demographics

Patient samples were classified according to ISSVA guidelines by a pathologist specialized in vascular anomalies (**Tables 1–2**). 35 patients were included in the study having VM (n=15) or AST (n=20). Additionally, 3 patients were classified as VM/AST having characteristic features of both vascular anomalies. All lesions were unifocal, except 3 multifocal VM lesions. Median age of the AST and VM patients were 18 (range 11-46 years, female-to-male ratio 14:6) and 31 (range 9-77 years female-to-male ratio 6:9), respectively. Most of the AST lesions (85%) located in the extremities. Of all AST lesions, 60% were intramuscular lesions (12/20), 5 lesions affected both intramuscular and subcutaneous tissue, and 1 both synovial membrane and intramuscular tissue. 60% of the VM lesions were from the extremities. Of all VM lesions, 3 were intramuscular, 2 located in both intramuscular and subcutaneous tissue, and 2 affected synovial membrane, intramuscular and subcutaneous tissue. All VM/AST lesions (n=3, 100%) were from the extremities, of which 1 was intramuscular. Genetic mutations were detected from vascular lesions by droplet digital PCR (ddPCR). Oncogenic PIK3CA variants were detected in 19/38 patients (AST 75%, VM 18%; **Table 1–2**), of which PIK3CA p.H1047R/L somatic mutation was found in 10/19 patients. 53% of patients had received sclerotherapy (VM 3/15, AST 15/20, VM/AST 2/3 patients, respectively). A representative magnetic resonance image of AST lesion locating in an ankle of a 13-year-old male is presented in **Fig. 1A**. A representative 3D confocal image of vessel organization in the AST lesion is shown in **Fig. 1B**. CD31-labelled longitudinal vessels were shown to be torturous, branched and variable in size (**Fig. 1B**).

**Figure 1.**
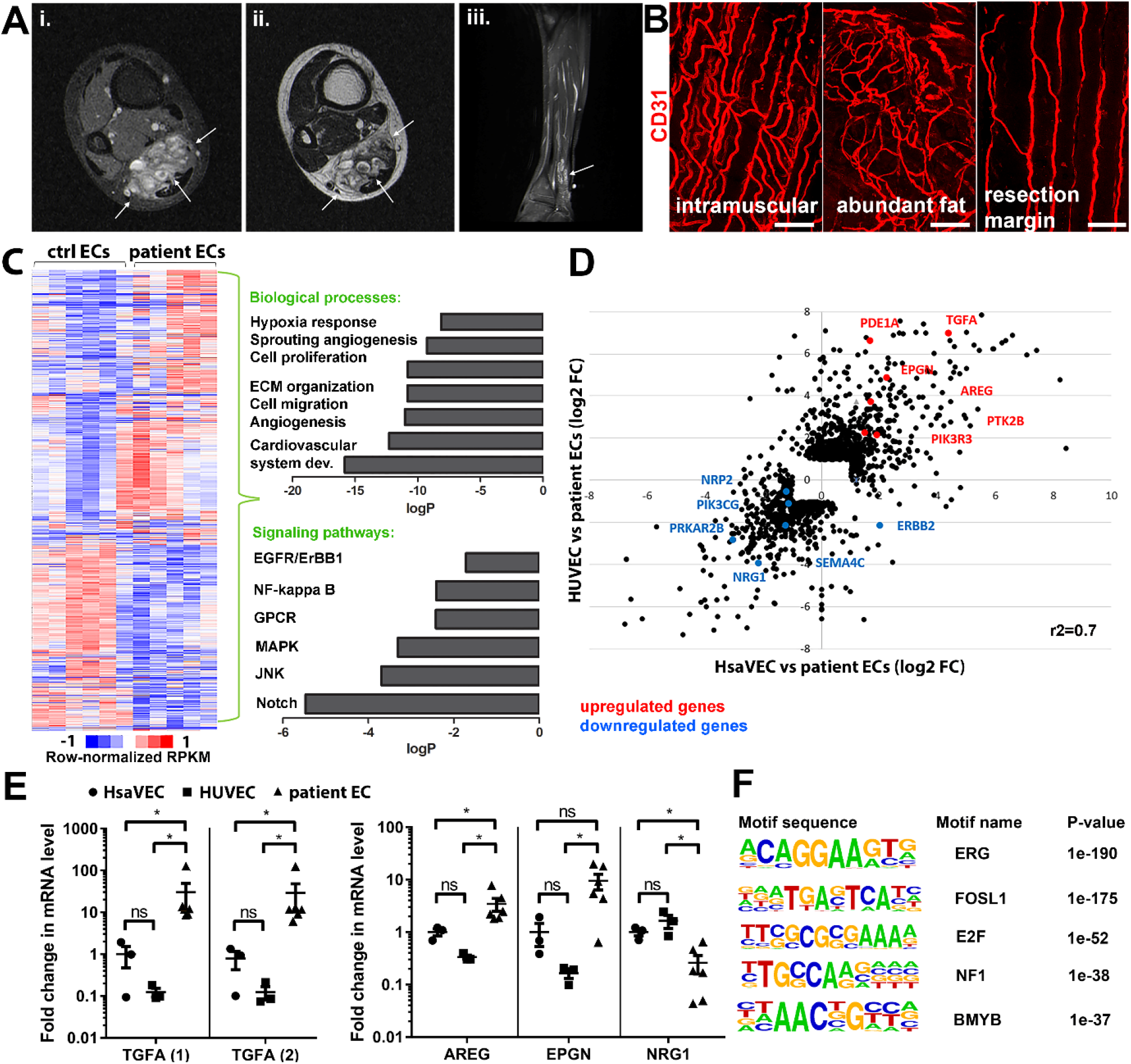
Genes involved in ErbB signaling pathway are upregulated in patient-derived ECs in vascular lesions with venous component. **A)** Magnetic resonance images of an AST lesion (arrows) in soleus muscle show the replacement of the normal muscle by dilated venous channels, diffusely enhancing small vessels and adipose tissue. i) Axial T1-weighted fat-saturated contrast enhanced image. ii) Axial T2-weighted image. iii) Sagittal T2-weighted fat-saturated image. **B)** 1mm-thick whole immunomounts were prepared from patient lesions, immunolabelled and imaged by laser scanning confocal microscopy. Images of the vasculature in AST lesion located in shin of a 16-year-old female are shown. Endothelial cells are immunolabeled with CD31 antibody (red). Vasculature of the same lesion in the intramuscular area (i), with abundant fat (ii) and next to resection margin (iii) are presented. Longitudinal vessels are seen. Scale bars, 100µm. **C)** Heatmap of normalized RPKM values (−1 to 1) of the differentially regulated genes in patient-derived ECs compared to HUVEC and HsaVEC control cells detected by bulk RNA-seq. Clustering was performed using Spearman’s rank correlation. Biological processes and cell signaling pathways detected by gene ontology analysis in patient-derived ECs. **D)** Scatter plot of the fold changes in gene expression comparing patient-derived and control ECs. Selected genes involved in PIK3CA, VEGFR2 and ErbB1-4 signaling are highlighted in red (upregulated) and blue (downregulated). Pearson correlation value (*r*^*2*^) is shown. **E)** Changes in mRNA expression levels of ErBB ligands (TGFA, with two different assays; amphiregulin, AREG; epigen, EPGN1; neuregulin 1, NRG1) were validated with RT-qPCR from patient-derived and control ECs. Mean and SEM are presented (HsaVEC and HUVEC, n=3; patient ECs, n=5). *, p < 0.05. **F)** Sequence motifs associated with differentially regulated genes in patient-derived ECs.

**Table 1.** Demographics of patients with AST patients.

| Patient | Gender | Age | Pathological diagnosis | # of lesions | Tissue <sup>a</sup> | Location | Somatic mutation (Fractional abundance <sup>d</sup>) |
| --- | --- | --- | --- | --- | --- | --- | --- |
| 1 <sup>b,c</sup> | F | 11 | AST | 1 | im | calf | - |
| 2 <sup>b,c</sup> | M | 34 | AST | 1 | im | shoulder | PI3KCA p.E542K (WT: 10.25; EC: 50.65; SC: none) |
| 3 <sup>b,c</sup> | M | 16 | AST | 1 | im | calf | PI3KCA p.H1047R (WT: 18.80; EC: 44.00; SC: none) |
| 4 <sup>b,c</sup> | F | 16 | AST | 1 | sc | shin | PI3KCA p.H1047L (WT: 8.30; EC: 48.95; SC: none) |
| 5 <sup>b</sup> | F | 17 | AST | 1 | im | back | - |
| 6 | M | 34 | AST | 1 | im | thigh | - |
| 7 | M | 17 | AST | 1 | im | thigh | PI3KCA p.E542K (WT: 7.32) |
| 8 | F | 31 | AST | 1 | im | thigh | PI3KCA p.H1047R (WT: 5.07) |
| 9 | F | 22 | AST | 1 | im | thigh | PI3KCA p.H1047R (WT 13.10) |
| 10 | F | 13 | AST | 1 | im, sc | foot | PI3KCA p.E545K (WT: 11.65) |
| 11 | M | 19 | AST | 1 | im | thigh | PI3KCA p.H1047R (WT: 8.89) |
| 12 | M | 13 | AST | 1 | im, sc | ankle | PI3KCA p.H1047R (WT: 12.60) |
| 13 | F | 23 | AST | 1 | im | calf | PIK3CA p.Y644H <sup>e</sup> |
| 14 | F | 41 | AST | 1 | sc | shin | - |
| 15 | F | 25 | AST | 1 | im | foot | PI3KCA p.E545K (WT: 8.93) |
| 16 | F | 16 | AST | 1 | im, sc | ankle | PI3KCA p.H1047R (WT: 5.98) |
| 17 | F | 46 | AST | 1 | im, sc | back | PI3KCA p.E542K (WT: 9.83) |
| 18 | F | 18 | AST | 1 | im, sc | ankle | PI3KCA p.E545K (WT: 11.35) |
| 19 | F | 13 | AST | 1 | im | calf | - |
| 20 | F | 24 | AST | 1 | im, sm | thigh | PI3KCA p.E542K (WT: 5.64) |

<sup>a</sup> Tissue: im, intramuscular; sc, subcutaneous; sm, synovial membrane.
<sup>b</sup> ECs and SCs isolated for cell experiments; <sup>c</sup> used in RNA-seq experiment.
<sup>d</sup> Fractional abundance of the mutation in WT, whole tissue lysate; EC, endothelial cells; SC, intervascular stromal cells; <sup>e</sup> mutation detected by whole-exome sequencing.

**Table 2.** Demographics of patients with VM.

| Patient | Gender | Age | Pathological diagnosis | # of lesions | Tissue <sup>a</sup> | Location | Somatic mutation (Fractional abundance in whole tissue lysate) |
| --- | --- | --- | --- | --- | --- | --- | --- |
| 21 | M | 34 | VM | 1 | im | thigh | TEK p.Y1108X <sup>d</sup> |
| 22 <sup>b,c</sup> | F | 77 | VM | 6 | sc | neck, fossa cubitalis, chest, hip, big toe | TEK p.L914F (5.63) |
| 23 | M | 40 | VM | 2 | im | chest, back | - |
| 24 | F | 69 | VM | 3 | im, sc | forearm <sup>e</sup>, hand | TEK p.L914F (10.01) |
| 25 | M | 24 | VM | 2 | sc | ankle, sole | PI3KCA p.H1047L (3.83) |
| 26 | M | 14 | VM | 1 | sc | lip | TEK p.L914F (16.31) |
| 27 | F | 39 | VM | 1 | im | chest | - |
| 28 | M | 28 | VM | 1 | sc | clavicle | TEK p.L914F (5.95) |
| 29 | M | 46 | VM | 1 | sc | shin | - |
| 30 | M | 31 | VM | 1 | sc | knee | TEK p.L914F (10.60) |
| 31 | F | 21 | VM | 1 | im, sc, sm | knee | - |
| 32 | M | 16 | VM | 1 | im, sc, sm | thigh, knee | - |
| 33 | M | 9 | VM | 1 | sc, sm | knee | - |
| 34 | F | 35 | VM | 1 | im, sc | blade | TEK p.L914F (12.10) |
| 35 | F | 16 | VM | 1 | sc | ankle | PI3KCA p.E545K (4.34) |
| 36 | M | 21 | VM/AST | 1 | im | upper arm | PI3KCA p.H1047R (3.91) |
| 37 | F | 41 | VM/AST | 1 | sc | ankle | KRAS p.Q61R <sup>d</sup> |
| 38 | F | 25 | VM/AST | 1 | sc | calf, leg | PI3KCA p.H1047L (5.23) |

119 <sup>a</sup> Tissue: im, intramuscular; sc, subcutaneous; sm, synovial membrane.120 <sup>b</sup> ECs and SCs isolated for cell experiments; <sup>c</sup> used in RNA-seq experiment.121 <sup>d</sup> mutation detected by whole-exome sequencing.122 <sup>e</sup> The patient had multiple lesions but only a lesion located in the forearm was operated.

### TGFA is upregulated in patient-derived ECs and vascular lesions with venous component

Five fresh tissue samples, including 4 AST and 1 VM, were obtained for bulk RNA-sequencing experiments. After digestion steps, ECs were selected by CD31 microbead kit. Somatic mutations in PIK3CA were detected by ddPCR in lesions from 3 out of 5 patients (**Table 1–2**), and for the first time, in ECs isolated from AST lesions (**Table 1**). Bulk RNA-sequencing was used to compare gene expression profiles of patient-derived^CD31+^ECs to healthy ECs derived from umbilical cord (HUVEC) or saphenous vein (HsaVEC). With principal component analysis, VM was not distinguished from AST samples and thus, was kept in the analysis. 1128 and 571 genes were found to be differentially expressed between control cells and patient-derived^CD31+^ECs, respectively (**Fig. 1C-D**). Differentially expressed genes (DEGs) were involved in angiogenic cellular processes, such as cell proliferation, migration, extracellular matrix organization and hypoxia (**Fig. 1C**; **Supplementary material 1**). Of particular interest were the fourteen genes found to be involved in ErbB signaling pathway known to regulate pathological angiogenesis. Multiple ligands of ErbB1-4 receptors were detected, e.g., transforming growth factor A (TGFA), amphiregulin (AREG), neuregulin-1 and epigen (EPGN). Also, G protein-coupled receptor signaling and RAS/MAPK cascade were found to be regulated **(Fig. 1C**; **Table 3**), previously linked to ErbB activation and downstream signaling (24,25). Similar genes and signaling pathways, e.g., TGFA and cell migration, proliferation, and ECM organization, were shown to be regulated in a separate analysis done for patient ECs with oncogenic PIK3CA variant only in comparison to control ECs (**Fig. 1 – figure supplement 1-2**).

Significant upregulation of ErbB1/EGFR ligands TGFA and amphiregulin (AREG) was validated by RT-qPCR in patient-derived^CD31+^ECs in comparison to control ECs, whereas no difference was observed in the regulation of ErbB4 ligand EPGN (**Fig. 1E**) (26,27). In accordance with data from bulk RNA-sequencing, ErbB2-ErbB4 binding neuregulin-1 was downregulated in patient-derived ^CD31+^ECs (**Fig. 1E**). De novo motif analysis of regulatory regions, i.e. enhancers within 100 kb of the gene transcriptional start site, was further used to identify possible regulatory transcription factor binding sites in patient-derived ECs that could regulate their phenotype. Cell cycle regulators E2F and FOSL1 were found to be the major regulators of transcription activity together with EC-specific transcription factor ERG (**Fig. 1F**).

**Table 3.** Selected cell signaling pathways regulated in patient-derived ECs.

| GO | Term | Genes |
| --- | --- | --- |
| GO:0038127 | ERBB signaling pathway | PRKCE,TNRC6C,PDE1A,**AREG**,<br>RPS27A,**TGFA**,KITLG,**ERBB2**,<br>PTK2B,DGKD,PRKAR2B,**NRG1**,<br>RPS6KA5,PRKACB |
| GO:0043122 | regulation of I-kappaB kinase/<br>NF-kappaB signaling | GREM1,LURAP1,BIRC3,TNFAIP3,<br>S100A4,MALT1,LITAF,PRKCE,<br>LPAR1,C18orf32,ZFAND6,RPS27A,<br>PLK2,TLR4,F2RL1,CASP1,S100A13 |
| GO:0008277 | regulation of G-protein coupled receptor<br>protein signaling pathway | RGS4,RGS9,DYNLT1,RGS11,CXCL8,RGS<br>10,RGS20,RGS5,RAMP2,RGS7,<br>PLCB1,ADRBK2 |
| GO:0043410 | positive regulation of MAPK cascade | PAK1,MAP4K2,SEMA4C,**TGFA**,<br>GADD45G,KSR2,**ERBB2**,NENF,<br>PLCB1,TPD52L1,GLIPR2,ICAM1,<br>CD74,PRKCE,LPAR1,PDCD10,INSR,<br>IGF1R,GADD45A,RPS27A,PTK2B,<br>KITLG,HGF,F2RL1,TLR4,PIK3CG,<br>ZEB2,**EPGN** |
| GO:0046328 | regulation of JNK cascade | PAK1,MAP4K2,GADD45A,IGF1R,<br>GADD45G,TNXB,PTK2B,CBS,SFRP1,PLC<br>B1,F2RL1,TLR4,TPD52L1,ZEB2 |
| GO:0007219 | Notch signaling pathway | HEY2,NOTCH1,TNRC6C,RBX1,<br>TMEM100,LFNG,RPS27A,E2F1,<br>DTX3,NOTCH2,MESP1,DNER,<br>FOXC1,SNAI2,HDAC9 |

\*ErbB pathway receptors and ligands are marked in bold.

**Figure 1 – figure supplement 1.**
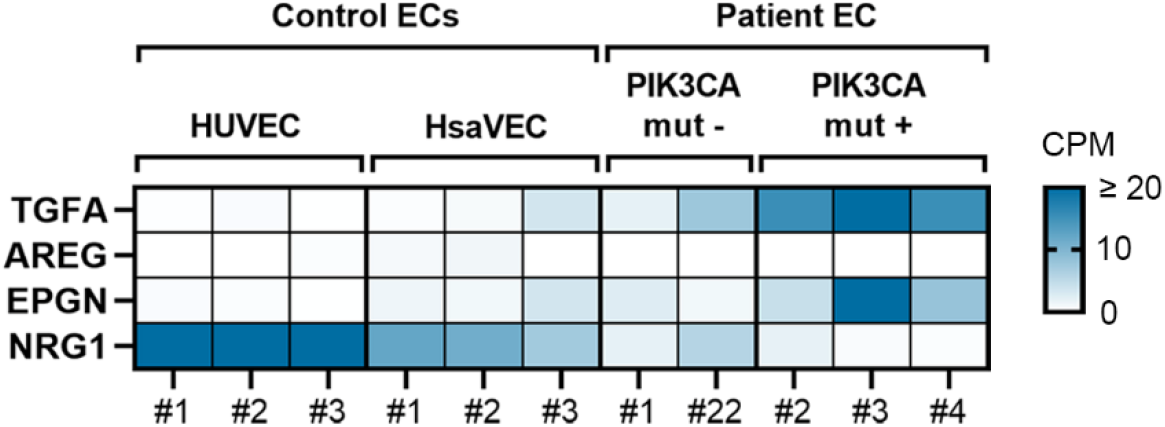
RNAseq data revealed that the highest levels of TGFA and EPGN mRNA were detected in patient-derived ECs having an oncogenic PIK3CA variant (PIK3CAmut+ Patient EC) in comparison to control ECs (HUVEC, HsaVEC) and patient-derived ECs without any oncogenic PIK3CA variant (PIK3CAmut-Patient ECs), while NRG1 mRNA was lower in patient-derived EC than control ECs. Data is presented in a heatmap format and shows normalized sequencing reads (counts per million reads, CPM) for the target gene expression in each sample separately. Scale 0-20 CPM (CPM values ≥ 20 are presented with the highest color intensity).

**Figure 1 – figure supplement 2.**
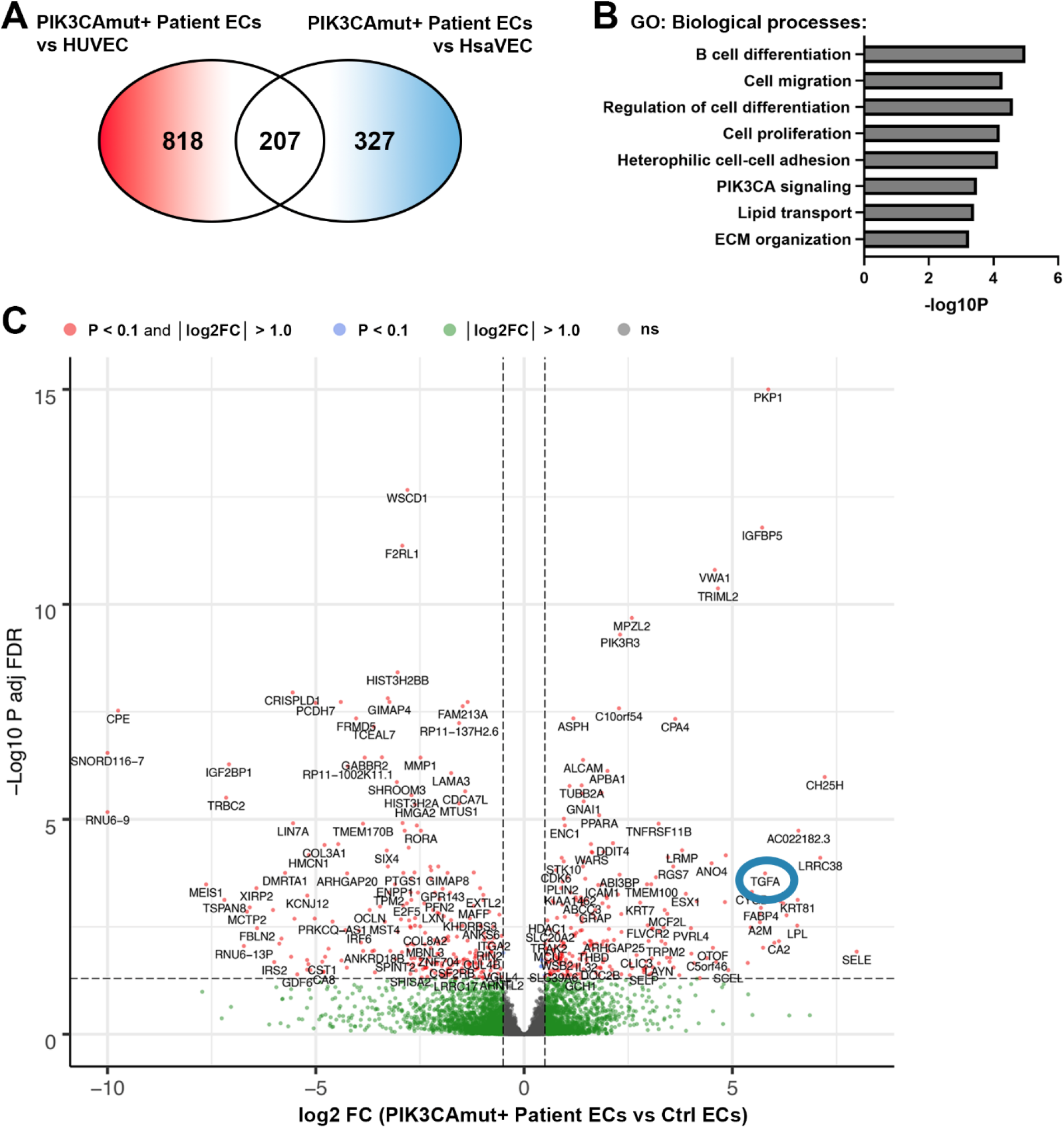
**A)** Further analysis demonstrated 818 DEGs between PIK3CAmut+ Patient ECs and HUVEC, and 327 DEGs between PIK3CAmut+ Patient ECs and HsaVECs (FDR-adjusted p-value < 0.1, log2 FC > 1.0 for both comparison). Since gene expression patterns in HUVECs and HsaVECs were rather similar (compared to patient-derived ECs), both control EC types were clustered together for the downstream analysis. Final analysis revealed 499 (FDR-adjusted p-value < 0.1, log2 FC > 1.0) DEGs between PIK3CAmut+ Patient ECs (n=3) and control ECs (HUVEC and HsaVEC, n=6). **B-C)** Biological processes detected by gene ontology analysis of DEGs **(B**) and a volcano plot of the genes **(C)** are presented. **C)** A blue circle highlights detection of TGFA as one of the top upregulated DEGs with connection to patient-derived ECs with an oncogenic PIK3CA variant.

Next, expressions of TGFA and AREG were validated at tissue level from patient lesions. TGFA mRNA was shown to be upregulated in both AST and VM by RT-qPCR (**Fig. 2A**, n=8 VM, n=3 AST) in comparison to control tissue, whereas no change of AREG mRNA was observed (**Fig. 2 – figure supplement 1A**). By immunohistochemistry, TGFA was detected in endothelium, pericytes and intervascular stromal cells in both AST and VM lesions (**Fig. 2B-C**; **Fig. 2 – figure supplement 1B**; positivity in 4/5 VM, 9/10 AST, 1/1 VM/AST lesions). Activated EGFR pathway was further demonstrated in AST lesions by detecting phosphorylated EGFR (positivity in 9/9 AST lesions; **Fig. 2D**), with most of the signal located in intervascular stromal cells. A heatmap for scoring of TGFA and pEGFR expression levels and the presence of oncogenic PIK3CA variant is presented in **Fig. 2E**, showing moderate or strong expression of these factors in the lesions.

**Figure 2.**
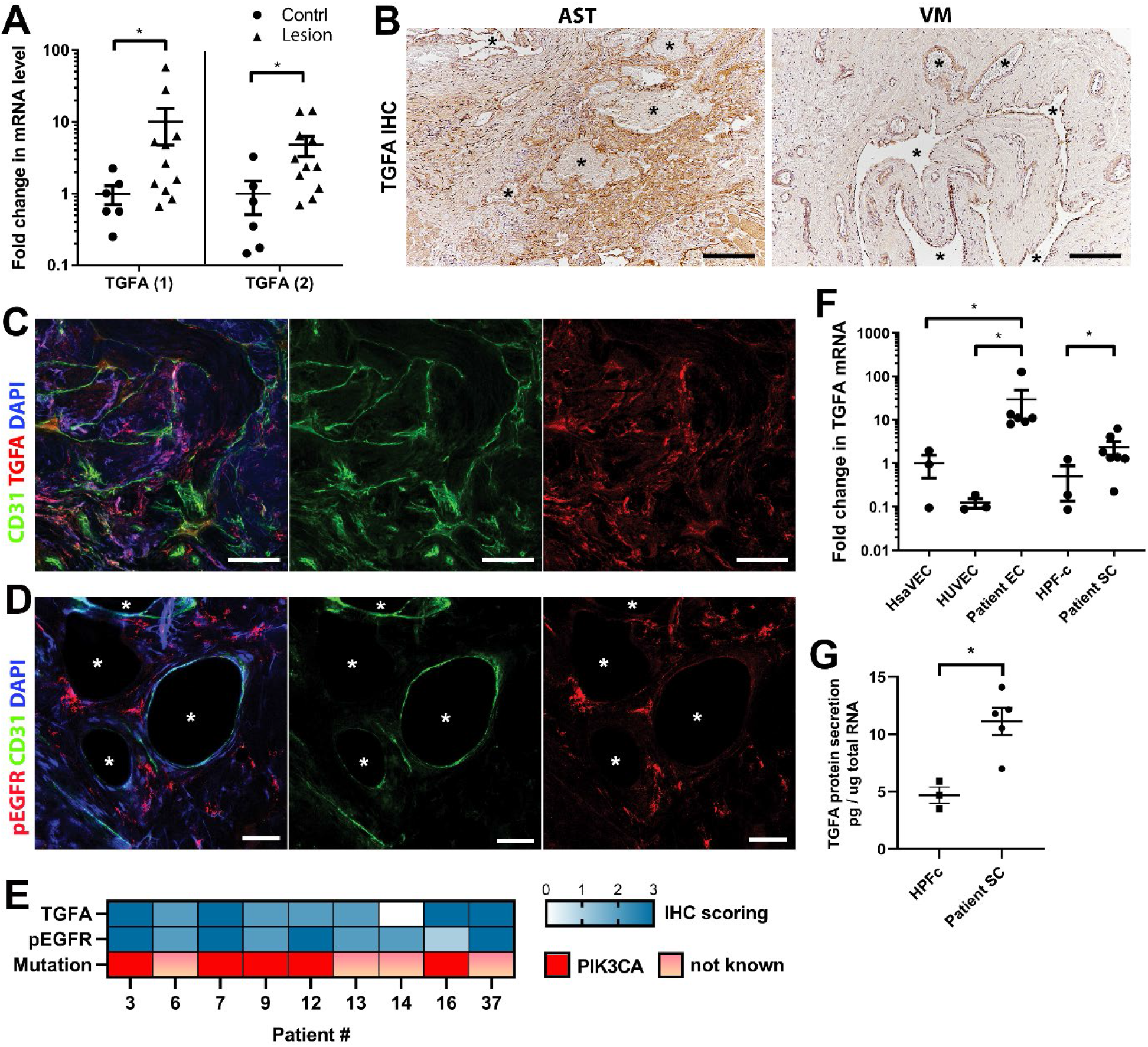
EGFR/ErbB1 ligand TGFA is upregulated in VM and AST patient tissue samples. **A)** RT-qPCR analysis showed significantly higher expression of TGFA mRNA (with two different assays) in VM and AST tissue than in control group. Mean and SEM are presented (lesions n=11; control group, n=6). *, p < 0.05. **B)** Representative images of TGFA expression in AST and VM patient samples. See Supplemental Figure 2B for normal skeletal muscle control. TGFA signal was detected in lesions in endothelium, pericytes and intrastromal cells by immunohistochemistry. Asterisks point out the largest vascular lumens which, especially in AST, are commonly tightly packed with erythrocytes. Scale bars, 200 µm. **C)** Representative whole immunomount images of CD31-labelled endothelium (green) and TGFA expression (red) in a patient diagnosed with intramuscular AST. Nuclei are stained with DAPI (blue). Longitudinal vessels are seen. Scale bars, 100µm. **D)** Representative confocal images of phosphorylated EGFR (red) expression in AST lesion. Endothelium is labelled with CD31 antibody (green), nuclei with DAPI. Vascular lumens are indicated with white asterisks in the cross-sections. Scale bars, 50µm. **E)** Heatmap of TGFA and pEGFR protein expression and presence of oncogenic PIK3CA variants in AST patients. Level of protein expression was scored (0-3) based on the detected signal in immunocytochemistry (0, none; 1, low; 2, medium; and 3, high). **F)** RT-qPCR analyses of TGFA expression in patient-derived ECs and intervascular stromal cell (SCs). Selection of ECs was performed by CD31 MicroBead Kit. Stromal cells were characterized by western blot and showed to be negative for EC marker CD31, and positive for fibroblast and smooth muscle cell marker vimentin (see Suppl. Fig. 2). The data is presented as relative mean fold change to HsaVEC control group and SEM (HsaVEC, HUVEC and HPF-c, n=3; patient ECs, n=5; patient SCs, n=6). *, p < 0.05. **G)** TGFA was shown to be secreted from patient-derived intervascular stromal cell (SCs) by ELISA (patient SC n=5; HPF-c n=3). **, p < 0.005.

In support of findings in immunohistochemistry, TGFA mRNA upregulation or secretion was detected in patient-derived^CD31+^ECs and in intervascular stromal^CD31-, vimentin+^ cells (Patient SCs; **Fig. 2F-G)**. By morphology patient SCs resembled fibroblasts, and (**Fig. 2F**) were characterized by western blot to be negative for CD31 marker and positive for fibroblast and/or smooth muscle cell marker vimentin (**Fig. 2 – figure supplement 1D-F**). None of the patient SCs had PIK3CA mutations detected by ddPCR (**Table 1**). Higher expression of EGFR mRNA was detected in control fibroblasts and patient SCs in comparison to ECs (**Fig. 2 – figure supplement 1C**). The data was in-line with scRNAseq data from mouse lower limb skeletal muscle (Tabula Muris, czbiohub.org) where Egfr did express in mesenchymal stem cells and skeletal muscle satellite cells but only in a small portion of ECs (**Fig. 2 – figure supplement 2A-B**). On the contrary, a small number of Tgfa^+^ cells were detected in mouse normal healthy skeletal muscle, showing the highest number of Tgfa^+^ cells in the EC cluster (**Fig. 2 – figure supplement 2C-D**).

Altogether, these results suggest that ErbB binding ligands are upregulated in AST and VM, and that TGFA, demonstrated to induce angiogenesis in earlier studies (26,28,29) could be among potential factors to induce a pro-angiogenic phenotype of lesion ECs.

**Figure 2 – figure supplement 1.**
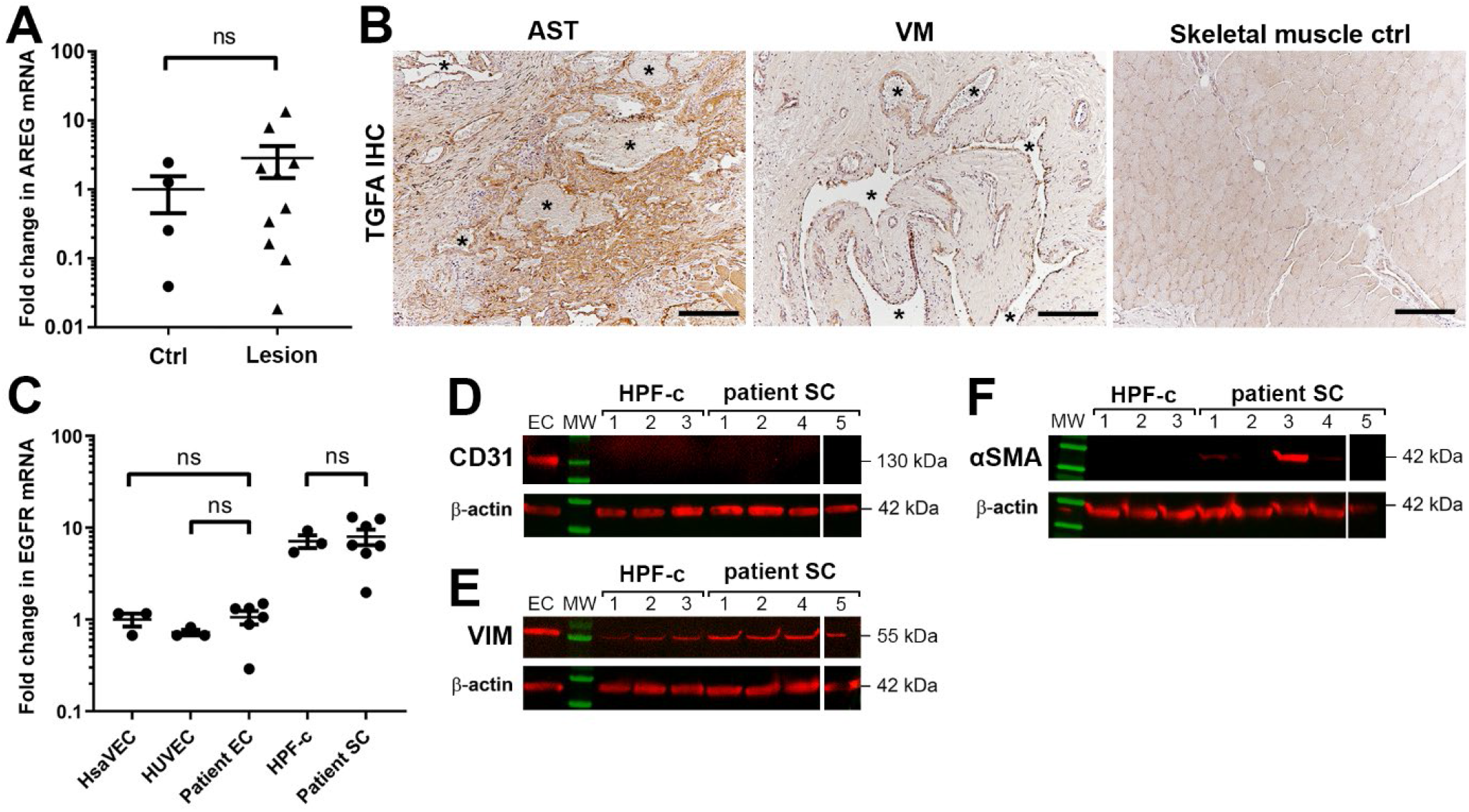
A) Expression of ErbB1 ligand amphiregulin in VM and AST lesions. RT-qPCR analysis of amphiregulin (AREG) mRNA in VM and AST lesions. Mean and SEM are presented (lesions n=10; control group, n=4; 1 lesion sample and 2 controls are not included in the blot due to having AREG mRNA expression under the detection limit). *, p < 0.05. **B)** Representative images of TGFA expression in AST and VM patient samples and in normal skeletal muscle control. TGFA signal was detected in lesions in endothelium, pericytes and intrastromal cells by immunohistochemistry. Asterisks point out the largest vascular lumens detected in the pathological samples. Scale bars, 200 µm. **C)** EGFR expression levels in patient-derived ECs and intervascular stromal cells. The data is presented as relative mean fold change to HsaVEC control group and SEM (HsaVEC, HUVEC and HPF-c, n=3; patient ECs and patient SCs, n=6). *, p < 0.05; ns, no significant difference. **D-F)** Western blot analysis for cell-type specific markers. Besides evident cell morphology, patient-derived intervascular stromal cells were confirmed to be CD31 negative **(D)**. All samples were positive for a fibroblast marker vimentin **(E)**. Some intervascular stromal cells were shown to be positive for aSMA **(F)**, a typical marker of activated fibroblasts and smooth muscle cells. **D-F)** β-actin was used as a control to confirm equal loading of the samples.

**Figure 2 – figure supplement 2.**
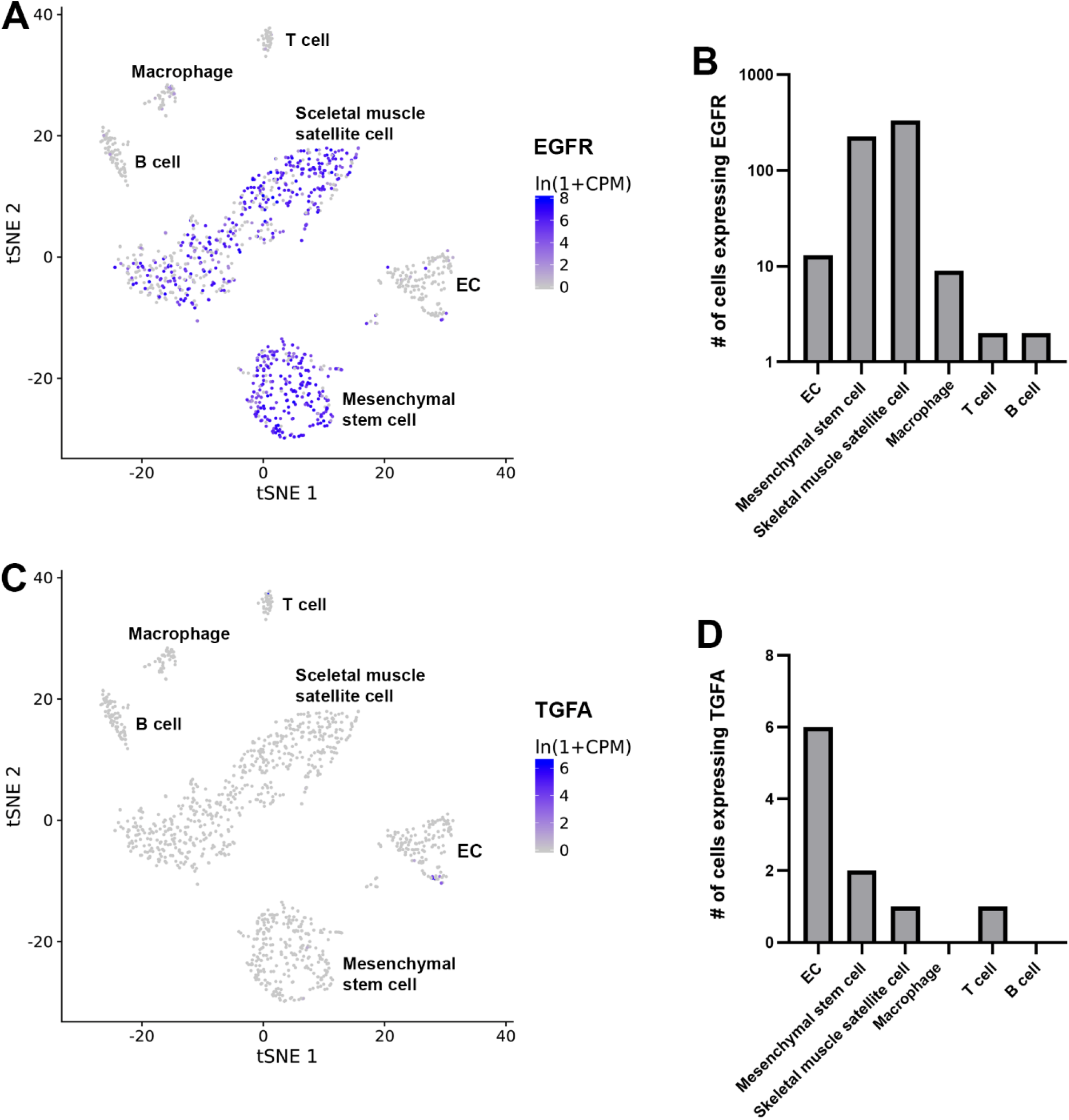
scRNAseq data from mouse lower limb skeletal muscle (Tabula Muris, czbiohub.org) of Egfr (**A-B**) and Tgfa (**C-D**) expression. Egfr was expressed mainly in mesenchymal stem cells and skeletal muscle satellite cells but only a small portion in ECs. A very few Tgfa^+^ cells were detected in mouse normal healthy skeletal muscle

### oncogenic PIK3CA p.H1047R induces expression of TGFA and enrichment of hallmark hypoxia

To understand the mechanism behind TGFA upregulation in patient lesions, and the potential role of oncogenic PIK3CA variant in it, bulk RNA-sequencing was performed on lentiviral-transduced ECs that expressed either wild-type or oncogenic PIK3CA p.H1047R, the most common somatic mutation found from this patient cohort (**Table 1–2; Fig. 3**; **Fig. 3 – figure supplement 1-2**). In-line with our experiments with patient-derived^CD31+^ECs, TGFA mRNA expression was shown to be induced in PIK3CA^H1047R^ expressing ECs (**Fig. 3 – figure supplement 1D**). In addition, GO analysis of these cells showed a hallmark “mTORC1 signaling”, indicative for activation of signaling pathway downstream from PIK3CA, as well as hallmarks “Glycolysis”, further indicative for a metabolic change from normal oxygen consumption towards anaerobic energy metabolism (**Fig. 3A)**. Interestingly, despite the normoxic cell culture conditions, hallmark hypoxia was detected as one of the top enriched hallmarks in PIK3CA^H1047R^ expressing ECs by GO analysis (**Fig. 3A**). Further comparison to the RNA-sequencing data from ECs expressing constitutively active hypoxia inducible factors (HIFs; GSE98060) confirmed that among 47 independent DEGs detected from our patient-derived^CD31+^ECs (**Fig. 1**) under “Response to hypoxia” (GO: 0001666) or “Hallmark Hypoxia”, majority (28 DEGs) were significantly altered by HIF1A, HIF2A or oncogenic PIK3CA p.H1047R. This indicated that part of the HIF regulated genes were also direct transcriptional targets downstream the oncogenic PIK3CA signaling. Top 15 hypoxia related DEGs are presented in **Fig. 3B**.

**Figure 3.**
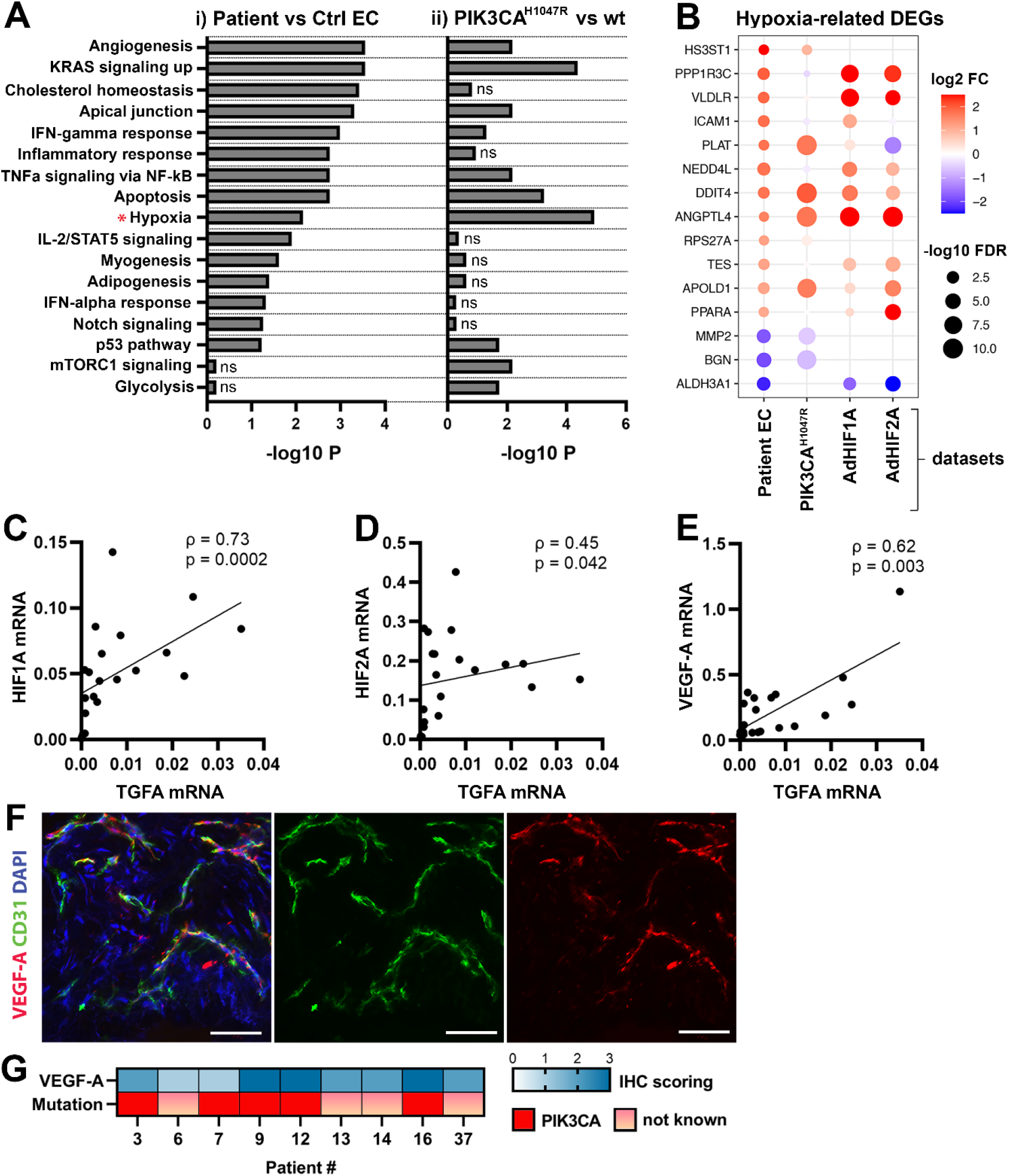
Oncogenic PIK3CA p.H1047R induces enrichment of hallmark hypoxia. **A)** Several shared MSigDB Hallmarks were found in bulk RNA-seq data from i) patient-derived ECs vs control ECs (left panel), and ii) ECs expressing PIK3CA wild-type (wt) or most common oncogenic variant detected in patient lesions, PIK3CA p.H1047R (right panel). Hallmark analysis was performed with the EnrichR web server, using adjusted p-value < 0.1 to define terms with significant enrichment of differentially expressed genes (DEGs). Hallmark Hypoxia (*) was detected as one of the top hallmarks in both RNA-seq datasets. **B)** Top 15 hypoxia-related genes differentially expressed between patient-derived vs control ECs, shown to be regulated in PIK3CA^H1047R^ expressing ECs and/or HIF1A/HIF2A. **C-E)** Correlation between TGFA and HIF1A (**C**), HIF2A (**D**) and VEGF-A (**E**) mRNA expression levels detected in AST and VM lesions (n=23). Spearman’s test was used to define correlation. Rho, Spearman’s rank correlation coefficient. **F)** Representative whole immunomount images of vasculature in AST lesion expressing VEGF-A (red) detected by confocal microscopy. Endothelium is labelled with CD31 antibody (green) and nuclei with DAPI (blue). Co-localization of VEGF-A and CD31 markers are seen in yellow. Longitudinal vessels are seen. Scale bars, 50 µm. **G)** Heatmap of VEGF-A expression and presence of oncogenic PIK3CA variants in AST patients. Level of VEGF-A expression was scored (0-3) based on the detected signal in immunocytochemistry (0, none; 1, low; 2, medium; and 3, high).

As HIFs and their target genes have previously been shown to induce angiogenesis or expression of TGFA and VEGF-A (30–33) and depletion of HIF1A/HIF2A have been shown to lead to downregulation in TGFA expression (34) correlation of these factors was next studied in lesions. The data showed a strong positive correlation between TGFA and HIF1A mRNA expression levels (ρ=0.73), and a moderate correlation between TGFA and HIF2A (ρ=0.45) (**Fig. 3C-D**, n=12 AST, n=7 VM, n=2 VM/AST). Additionally, a positive correlation (ρ=0.62) between VEGF-A, and TGFA mRNA expression levels (**Fig. 3E**) was observed. VEGF-A expression or secretion was further validated by immunohistochemistry in patient lesions (**Fig. 3F-G**) or in patient SCs by RT-qPCR and ELISA (**Fig. 3 – figure supplement 3**).

**Figure 3 – figure supplement 1.**
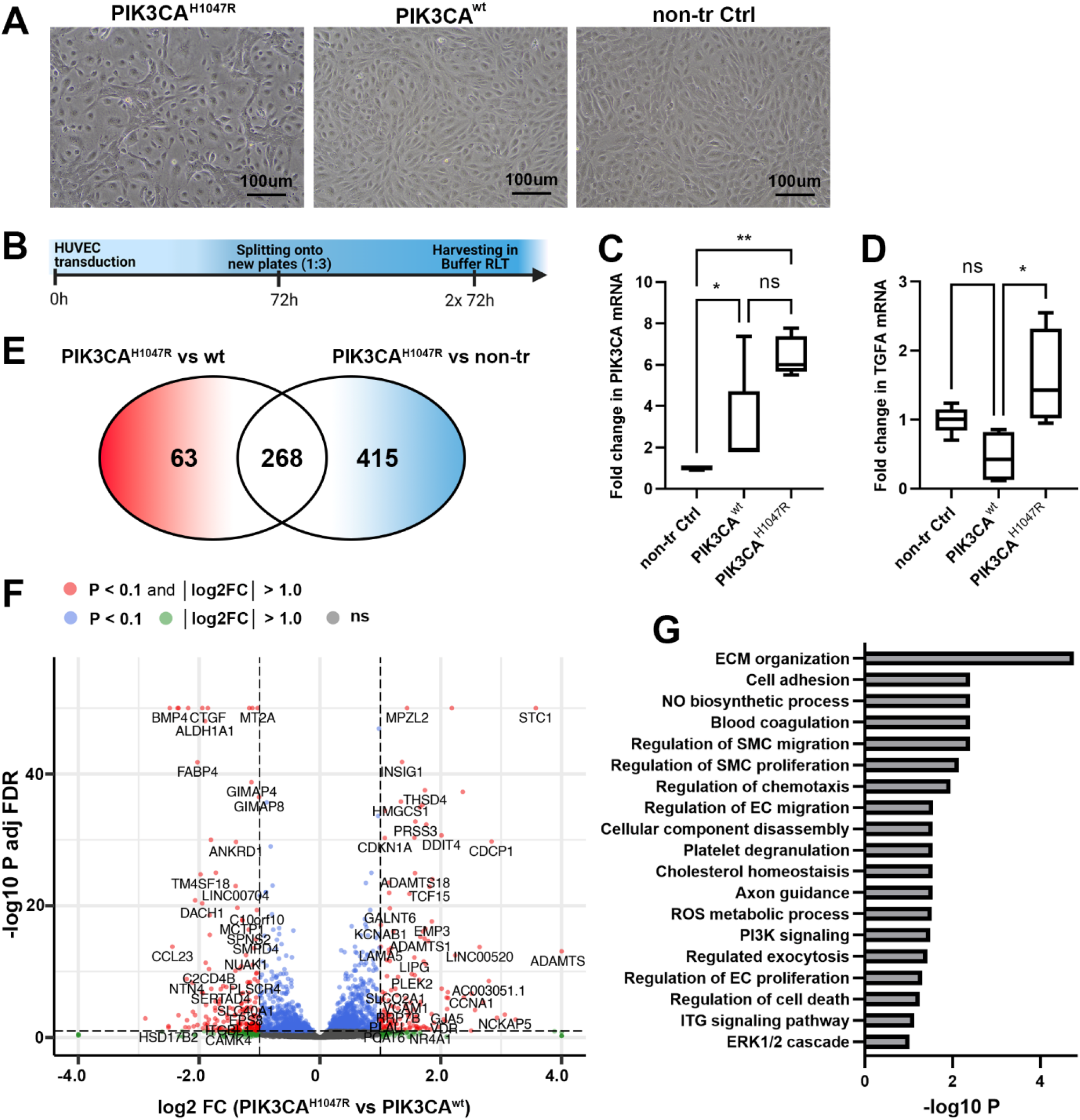
**A)** In-line with previous publications (9), expression of oncogenic PIK3CA variant p.H1047R induced morphological changes in HUVECs. Representative images of HUVECs transfected with lentiviral vector (LV) encoding PIK3CA^H1047R^ and PIK3CA^wt^ **(A)**, in comparison to morphology of non-transduced HUVECs. Scale bars, 100µm. **B)** Schematics showing preparation of cell culture samples for RNA-sequencing experiments to analyze transcriptional changes induced by PIK3CA^H1047R^ in HUVECs. PIK3CA^wt^-transduced and non-transduced HUVECs were used as controls (n=4 in each group). **C-D)** Significantly higher expression of PIK3CA mRNA was detected in cells expressing PIK3CA^H1047R^ or PIK3CA^wt^ than non-transduced cells, confirming that lentiviral transduction worked, and the vectors are functional (**C**), meanwhile TGFA mRNA was significantly induced in ECs expressing PIK3CA^H1047R^ **(D)**. Data is representative from two independent experiments done in 5-8 replicates and shown as mean and SEM. *, p < 0.05; **, p < 0.01). **E-G)** RNA-sequencing revealed 268 genes that were differently expressed in PIK3CA^H1047R^ than PIK3CA^wt^ or non-transduced HUVECs, demonstrated using Venn diagram **(E)** and Volcano plot **(F)**. Biological processes detected by gene ontology analysis of differentially expressed genes **(G)**.

**Figure 3 – figure supplement 2.**
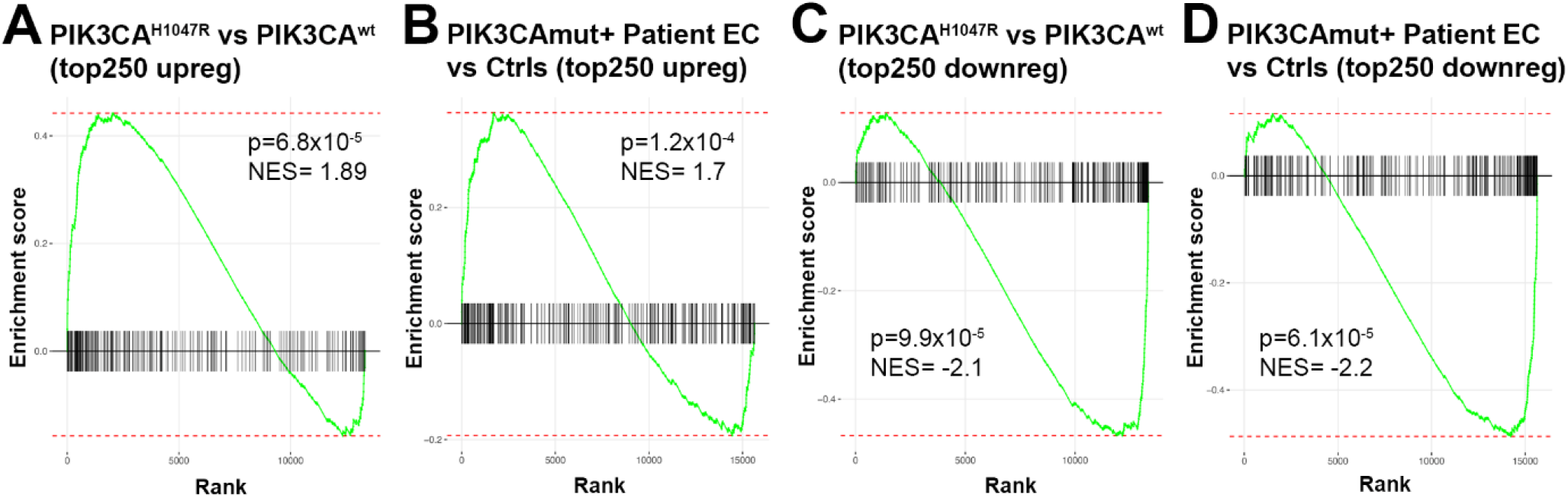
Gene set enrichment analysis (GSEA) done for upregulated **(A-B)** or down-regulated **(C-D)** DEGs confirms that the RNAseq data from HUVECs overexpressing PIK3CA^H1047R^ (vs PIK3CA^wt^) is well in-line with the RNAseq data from patient-derived ECs (vs control ECs); thus, justifying that PIK3CA^H1047R^ (vs PIK3CA^wt^)-transduced ECs is a feasible model to be used in further experiments. In the GSEA analysis, top 250 DEGs upregulated in PIK3CA^H1047R^ vs PIK3CA^wt^-transduced HUVECs were compared to the custom gene set consisting of top 250 DEGs upregulated in PIK3CAmut+ Patient ECs vs control ECs (HUVEC, HsaVEC; all EC types from 3 donor) **(A)** and vice versa **(B)**. Top 250 downregulated DEGs from both RNAseq experiments were compared similarly, by using data from PIK3CAmut+ patient ECs vs control ECs as a query set in **(C**) and data from PIK3CA^H1047R^ vs PIK3CA^wt^-transduced HUVECs as a query set in **(D)**. p-values between 6.1×10^−5^ and 1.2×10^−4^ are considered as a high significance for correlation between the data sets, NES, normalized enrichment score.

**Figure 3 – figure supplement 3.**
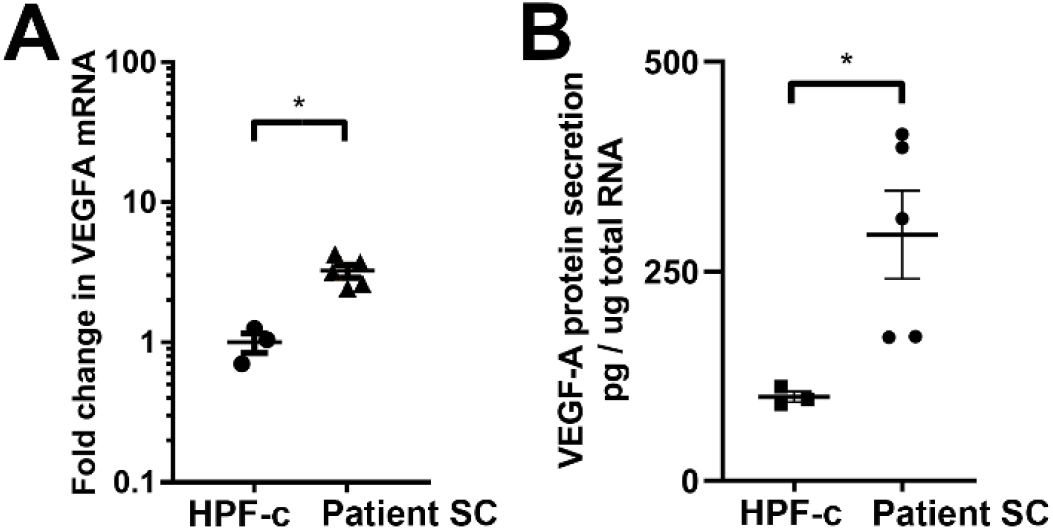
**A)** RT-qPCR analysis of VEGF-A mRNA expression in patient SCs and control fibroblasts (SCs, n=6; HPF-c, n=3). *, p < 0.05. **B)** VEGF secretion from patient SCs analyzed by ELISA (SC n=5; HPF-c n=3). *, p < 0.05.

### Patient SCs and TGFA induce a pro-angiogenic EC phenotype together with VEGF-A

To study further the effect of VEGF/TGFA-secreting patient SCs on ECs, we used a modified fibrin bead angiogenesis assay (35). Patient SCs were shown to induce HUVEC sprouting without any additional growth factor stimulation (**Fig. 4A-B**), implying that paracrine factors secreted by the SCs modulate the EC phenotype. We hypothesized that the mechanism could be via TGFA-mediated upregulation of VEGF-A. Accordingly, a significant increase in VEGF-A mRNA and protein secretion were observed after stimulation of control fibroblasts (HPF-c) with rhTGFA (**Fig. 4 – figure supplement 1)**. Also, in a fibrin bead assay with HUVECs, co-stimulation with rhVEGF-A and rhTGFA proteins resulted in a higher increase of EC area and sprouting in comparison to either of the growth factors alone (**Fig. 4C-E**). This suggested that these growth factors have a synergistic effect on modulating EC phenotype.

**Figure 4.**
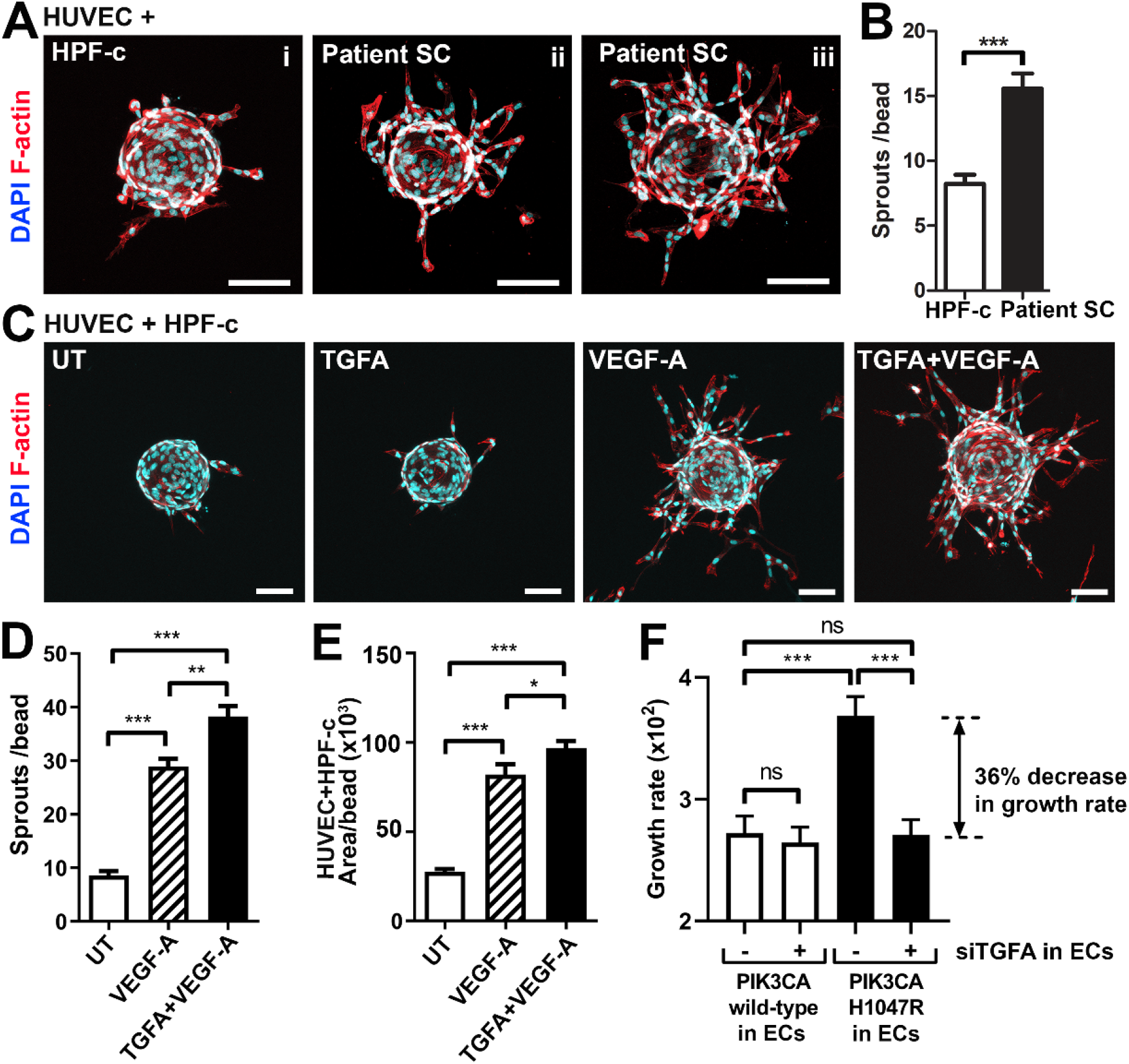
Patient SCs and TGFA induce an angiogenic EC phenotype together with VEGF-A. **A-B**) VM patient SCs induce sprouting of genotypically normal ECs. HUVECs on collagen-coated beads were embedded into a fibrin gel and patient SCs or control HPF-c cells were put on top. Representative images are presented at d7. ECs are labelled with phalloidin (red). nuclei with DAPI (blue; **A**) The number of sprouts per bead in each condition is shown **(B)**. 2 independent experiments were done in triplicates. *, p < 0.05. In all images, scale bar 100µm. **C-E)** Fibrin bead assay with HUVECs and HPF-c cells shows increased EC sprouting after stimulation with rhVEGF-A and rhTGFA at d6. ECs are labelled with phalloidin (red), nuclei with DAPI (blue; **C**). The number of sprouts per bead **(D)** or sprout area **(E)** in each condition was determined from confocal images by ImageJ (45 beads/group). 2 independent experiments were done in triplicates. ***, p < 0.001. **F)** Co-culture experiments with HPF-c and HUVEC cells showed increased growth rate in wells with PIK3CA^H1047R^-expressing ECs in comparison to wells with ECs expressing PIK3CA^wt^. The response was abolished after inhibition of endogenous TGFA in ECs by specific siRNA, demonstrating the involvement of TGFA in PIK3CA^H1047R^-induced responses. Wells with siCtrl-transduced ECs (marked to be negative for siTGFA) were used as a control group in the experiments. Cellular growth was monitored using IncuCyte Live-Cell Imaging system. Data is presented as relative growth rate from 2 experiments done in triplicates. ***, p < 0.001. In all data, mean and SEM are presented.

In contrast to TGFA (**Fig. 3 – figure supplement 1D**), no change in VEGF-A mRNA expression was seen in PIK3CA^H1047R^ expressing ECs. Neither did rhTGFA significantly affect cell proliferation in cultures with ECs or HPF-c alone. To understand the importance of TGFA expression in the growth of ECs in the presence of PIK3CA^H1047R^ mutation and HPF-c or SC, proliferation assays were performed in co-culture conditions. Co-culturing of PIK3CA^H1047R^-expressing ECs together with HPF-c showed increased growth rate in comparison to PIK3CA^wt^-treated cells. Importantly, the response was abolished when TGFA expression in ECs was knocked down by siRNA (**Fig. 4F**; **Fig. 4 – figure supplement 2**). Thus, altogether the data suggests that TGFA expression, induced by oncogenic PIK3CA p.H1047R, results in a pro-angiogenic EC phenotype by increasing cell proliferation and VEGF-A secretion but only in conditions where ECs and HPF-c/SCs are combined.

**Figure 4 – figure supplement 1.**
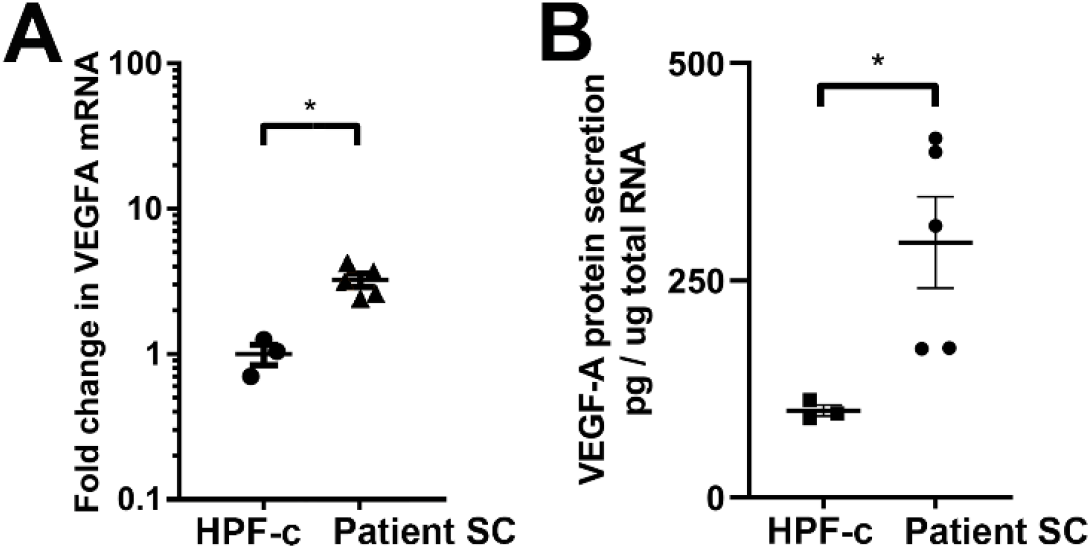
rhTGFA stimulation of HPF-c cells increased VEGF-A mRNA expression **(A, 6h)** and VEGF-A protein secretion **(B, 24h)**. The data is from three independent experiments done in triplicates. ***, p < 0.001.

**Figure 4 – figure supplement 2.**
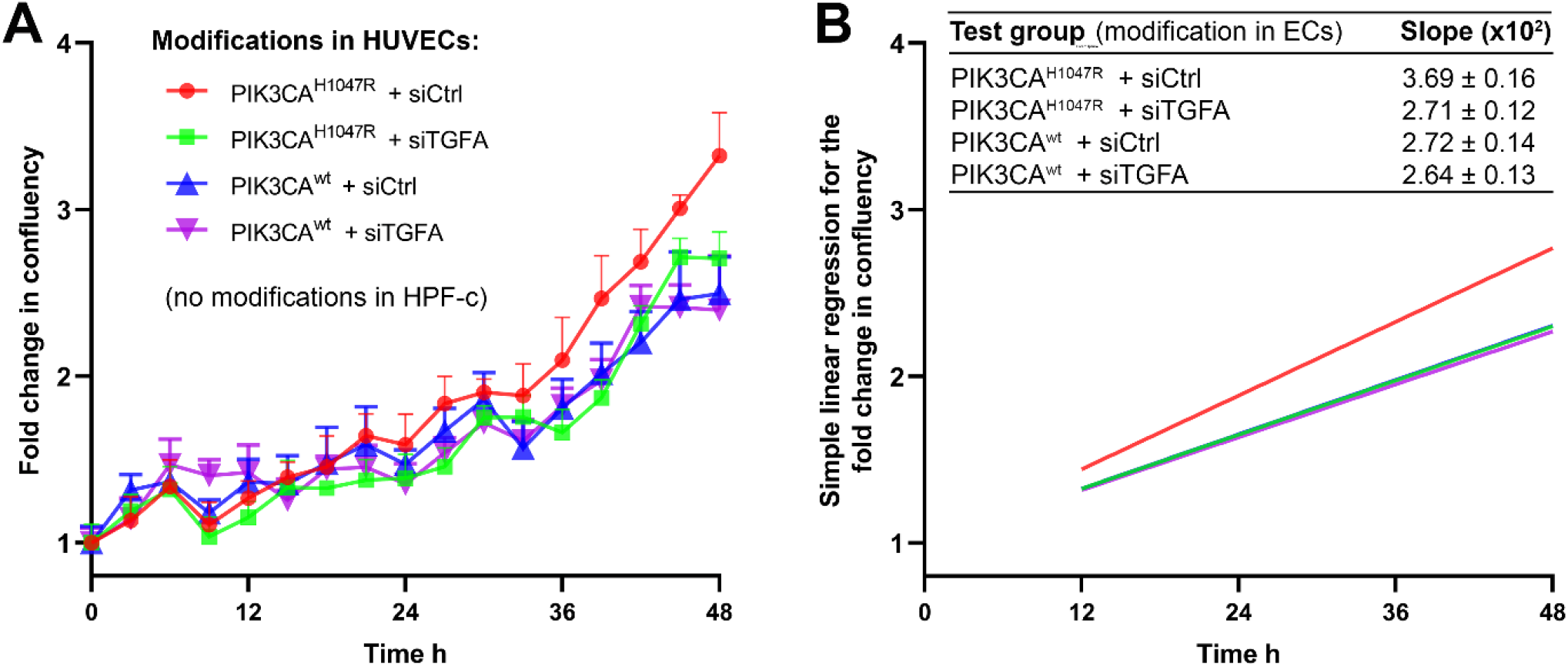
Quantification for proliferation of cells in co-culture experiments combining ECs expressing PI3KCA^H1047R^ or PI3KCA^wt^ +/-TGFA and genetically normal HPF-c. Knock-down of TGFA was performed using siRNA targeting to TGFA and non-targeting control siRNA as a control. Just prior imaging, HUVECs and HPF-c were mixed at ratio 8:1 to induce crosstalk between the cell types in cultures without external addition of growth factors. Cell growth was monitored using IncuCyte S3 Live-cell Imaging System for 48h in 3h intervals, 4 images/well. **A)** Growth curves for each condition show fold change in confluency of cells in relation to time. Data is presented as mean and SEM from 2 experiments done in triplicates. **B)** Simple linear regression for cell growth in each condition. Regression lines were generated between time points 12-48 h, as the first 12 hours were considered as a time when the cells settle on wells after seeding. Mean and SEM of slopes of the regression lines, indicating growth rate of the cells, are shown in the table.

### Fibroblasts induce vascularization in a mouse xenograft model for vascular lesion

Due to the small number of cells obtained from patient lesions, further studies to understand the role of SCs/fibroblasts in lesion formation in vivo were performed with commercially available primary cells. A new modified mouse xenograft model based on (36) was used for the first time combining: i) ECs expressing either oncogenic PIK3CA p.H1047R or PIK3CA wild-type, and ii) genotypically normal primary fibroblasts (HPF-c; **Fig. 5A**). Lesion growth and size at d20 was observed to be similar between PIK3CA^H1047R^-transfected ECs with or without fibroblasts (**Fig. 5B**). With H&E staining, various sized vascular channels filled with erythrocytes were detected **(Fig. 5D**). Notably, there was no blood-filled vascularization shown in explants with ECs expressing PIK3CA^wt^, whereas vascular channels in explants with PIK3CA^wt^ expressing ECs and fibroblasts clearly contained erythrocytes (**Fig. 5C-D, 5H**). Higher vascularization, detected with EC marker CD31, was observed with explants with embedded fibroblasts in comparison to ECs alone (**Fig. 5E-F**). In comparison to explants containing only PIK3CA^H1047R^ ECs, the vascular channels with fibroblasts were wider and the endothelium appeared to be more organized **(Fig. 5E-F)**. The highest CD31 vascularization score was detected in explants with ECs expressing PIK3CA^H1047R^ and fibroblasts (**Fig. 5G)**. In addition, a higher number of inflammatory cells were seen in the explants with ECs expressing PIK3CA^H1047R^ than PIK3CA^wt^; however, no statistical difference was detected with or without fibroblasts (**Fig. 5I**).

**Figure 5.**
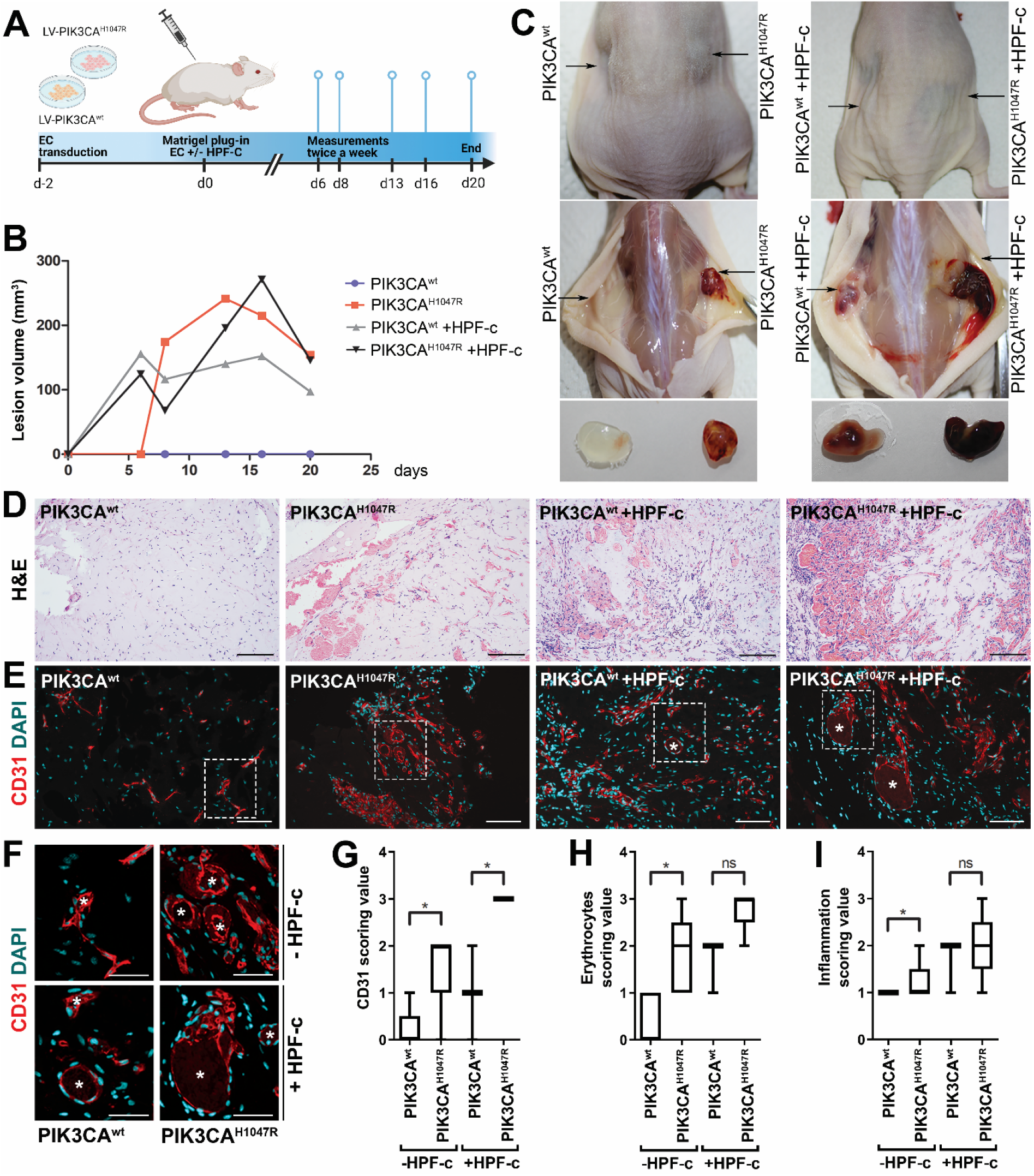
Fibroblasts induce vascularization in a mouse xenograft model for vascular lesion. **A)** Subcutaneous injection of matrigel with HUVECs transduced with PIK3CA^wt^ or PIK3CA^H1047R^ encoding lentivirus vectors, with or without primary fibroblasts, was performed in athymic Nude-Foxn1^nu^ mice. A timeline of the animal experiment is presented. **B)** Lesion volume measured by caliper from day 6 to day 20 (n=5 EC^wt^, n=5 EC^H1047R^, n=3 EC^wt^+FB, n=5 EC^H1047R^+FB). **C)** Representative images of mice and dissected lesion explants on day 20. **D, E)** Explant sections stained with hematoxylin and eosin (H&E; **D**) or EC marker CD31 (red) and DAPI (nuclei, blue; **E**). Scale bars, 200µm, H&E; 100µm, CD31. **F)** Close-up of the vascular lumens detected in the explants with or without fibroblasts (CD31, red; DAPI, blue). Scale bars, 50µm. **G**) Scoring for vascularization done for sections stained for CD31. The highest vascularization was observed in the explants with HUVECs expressing PIK3CA^H1047R^ and fibroblasts. *, p < 0.05. **H, I)** Scoring for erythrocytes **(H)** and inflammation **(I)** done on H&E-stained sections. *, p < 0.05.

Further immunohistochemistry confirmed production of EGFR protein in the explants with PIK3CA^H1047R^-expressing ECs; however, no difference was detected between explants with or without fibroblasts (**Fig. 5 – figure supplement 1**). In summary, the data indicates the importance/potential of fibroblasts in inducing aberrant vasculature in lesions.

**Figure 5 – figure supplement 1.**
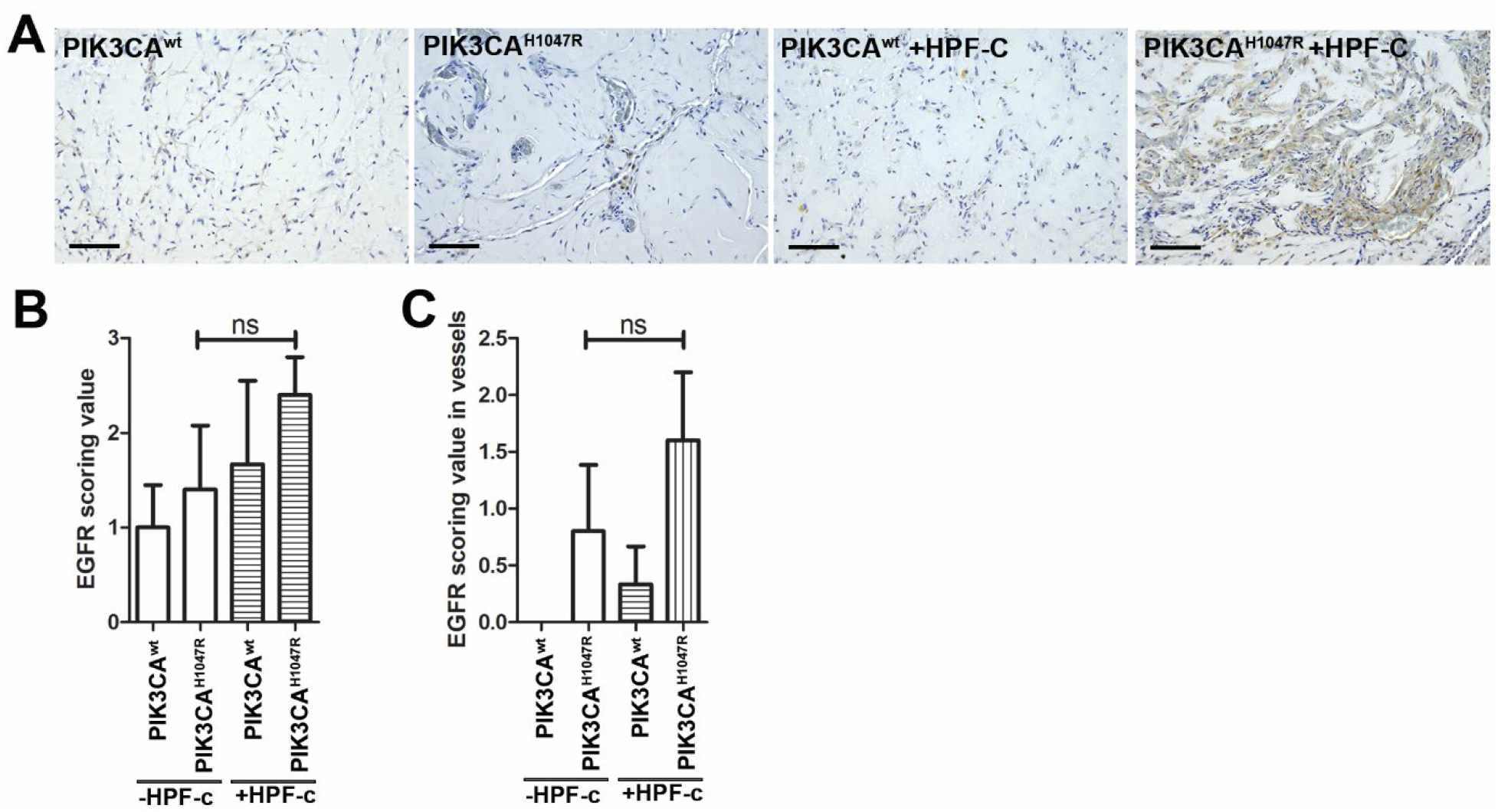
Subcutaneous injection of matrigel with HUVECs transfected with wt or PIK3CA^H1047R^ expressing lentivirus vectors with/without primary fibroblasts was performed in athymic Nude-Foxn1^nu^ mice. **A)** Explant sections stained with EGFR antibody. Scale bars, 100µm. **B-C)** EGFR expression was scored in the explants in the lesion area (**B**) and in vascular structures (**C**).

### ErbB family antagonist Afatinib reduces VEGF secretion, angiogenic sprouting and lesion size

Next, afatinib (Gilotrif™, Giotrif®), an inhibitor of EGFR/ErbB1, ErbB2 and ErbB4, was used to test the potential inhibition of EC angiogenesis and vascular lesion growth in mice. First, *in vitro*, afatinib was shown to block TGFA-stimulated EGFR phosphorylation detected by western blot (**Fig. 6A**), and to decrease VEGF-A secretion from TGFA-stimulated control fibroblasts measured by ELISA (**Fig. 6B**). Accordingly, in the fibrin bead assay, afatinib was shown to reduce rhVEGF-A and rhTGF-A mediated EC sprouting (**Fig. 6C-D**). To further test the effect of afatinib on PI3K-driven vascular lesion growth, our modified mouse xenograft model was used with ECs expressing oncogenic PIK3CA^H1047R^ and genotypically normal primary fibroblasts. Lesions were allowed to form for 9 days, reaching 200.1±10.2µm^3^ in size, followed by afatinib treatment daily p.o. for 9 days (**Fig. 7A**). At d18, lesion size (**Fig. 7B-C**) and vascularization detected by H&E and CD31 stainings (**Fig. 7H-L**) was shown to be reduced in afatinib-treated mice in comparison to untreated mice. In both groups, various sized vascular channels filled with erythrocytes were seen **(Fig. 7J-K**). A lower number of inflammatory cells was detected in the afatinib treatment group (**Fig. 7K**). Accordingly, reduction of EGFR expression was detected in explants of afatinib-treated mice (**Fig. 7I, 7M**). Altogether, the data validates that ErbB signaling plays a key role in PIK3CA p.H1047R lesion formation in the presence of fibroblasts and provides a new potential therapeutic strategy for targeting vascular lesions with a fibrous component.

**Figure 6.**
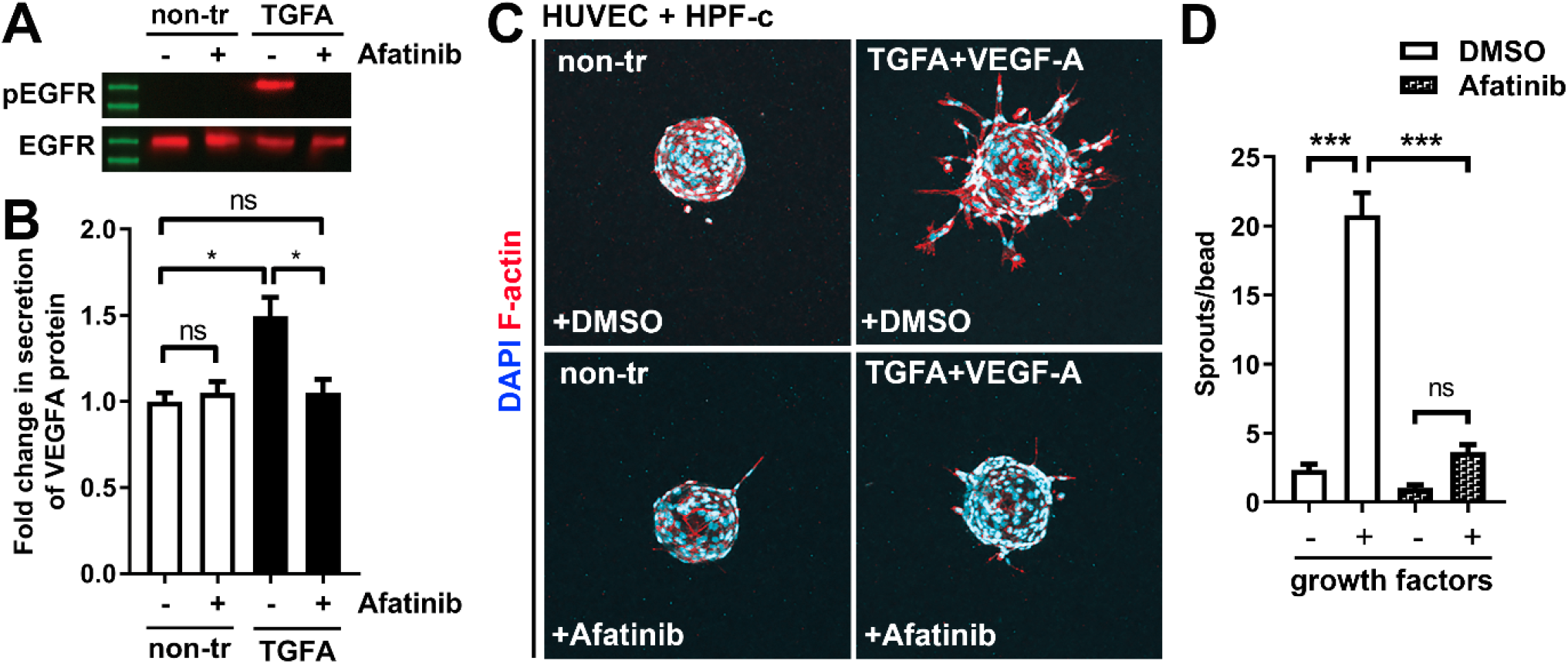
Afatinib reduces VEGF-A secretion, angiogenesis and lesion size. **A)** Afatinib decreased EGFR/ErbB1-phosphorylation in rhTGFA-stimulated HPF-c cells. Total EGFR was used to control equal loading of the samples. **B)** VEGF secretion measured by ELISA from rhTGFA-stimulated control fibroblasts (HPF-c) with or without afatinib treatment. Two independent experiments done in triplicates is presented as mean and SEM. **, p < 0.01. **C, D)** In the fibrin bead assay with HUVECs and HPF-c, afatinib inhibited EC sprouting induced by co-stimulation with rhVEGF-A and rhTGFA. Representative images of each group are presented at d7 **(C)** ECs are labelled with phalloidin (red) and nuclei with DAPI (blue). Quantitative analysis for the number of sprouts per bead **(D)** was performed with ImageJ software (30 beads/group). Afatinib treatment was started at d3 after HUVECs had already formed angiogenic sprouts. The data from two independent experiments done in triplicates is presented as mean and SEM. *, p < 0.05; ***, p < 0.001.

**Figure 7.**
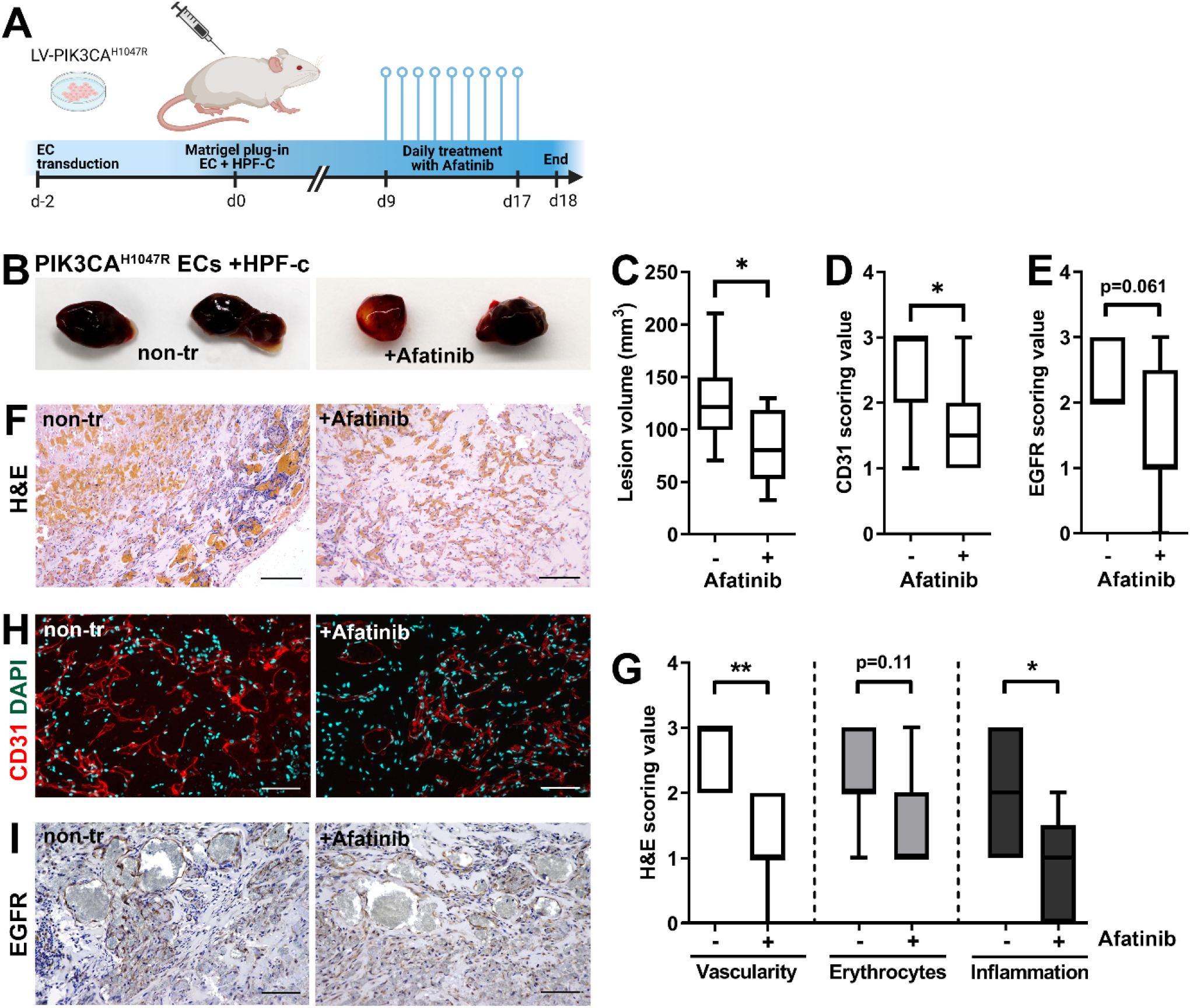
Afatinib reduces lesion size in matrigel plug-in assay. Subcutaneous injection of matrigel with HUVECs transfected with PIK3CA^H1047R^ expressing lentivirus vectors with primary fibroblasts was performed in athymic Nude-Foxn1^nu^ mice. After lesions reached 200µm^3^ in size, afatinib treatment was started for 9 days (25 mg/kg, p.o., daily). **A)** A timeline of the animal experiment is presented. **B)** Representative images of dissected explants on day 18. **C)** Lesion volume measured from dissected explants at d18 (n=7 untreated, n=9 afatinib treated). **D)** Scoring for vascularization done for sections stained for CD31 (**H**, n=7 untreated, n=9 afatinib treated). **E)** Scoring for EGFR expression (**I**; n=6 untreated, n=9 afatinib treated). **F**,**H)** Explant sections stained with hematoxylin and eosin (H&E; **F**) or EC marker CD31 (red) and DAPI (nuclei, blue; **H**). Scale bars, 200µm, H&E; 100µm, CD31. **G)** Scoring for vascularization, erythrocytes and inflammatory cells done on H&E-stained sections. **I)** Explant sections stained with EGFR. Scale bars, 100µm.

## DISCUSSION

Symptomatic AST and VM are primarily treated with compression garments, and if needed, with percutaneous sclerotherapy, percutaneous cryotherapy, endovascular laser treatment or surgical resection (14–18). In sclerotherapy, sclerosants are administrated using ultrasound guidance intravenously to induce endothelial damage that leads to a total or partial atrophy of the lesion. Single sclerotherapy rarely results in an adequate treatment response but often reduces the lesion size alleviating the symptoms (37). In our earlier study, insufficient response to sclerotherapy was detected primarily in patients with lower-extremity intramuscular AST lesions (5). Thus, surgery has been used as the primary treatment for AST, whereas percutaneous sclerotherapy is often the first treatment option for VM. Due to high recurrence, difficult anatomical location, possible functional impairment associated with operation, and a risk of tissue necrosis after sclerotherapy, more effective therapies are, however, needed for the treatment of AST and VM. Previously, mTOR inhibitor sirolimus has been tested in clinical trials with promising results for patients having VM and somatic mutations leading to constitutive activation of the PI3K pathway (19,20) (ClinicalTrials.gov, study nro: NCT02638389).

In sporadic VM, PI3K/Akt activating somatic TEK mutations are associated with skin lesions and p.L914F mutation is found in 60% of the patients (9,10). Genetics and disease mechanisms in non-skin associated VMs are less defined. VMs with somatic mutations in PIK3CA do not extend to skin (9) and are found in approximately 20% of the patients. In the study of Castel et al. (2016), 24% of the patients (4/17) having intramuscular sporadic VM lesions had mutation in PIK3CA gene and 2 out of 17 patients in TEK p.L914F (38). Prior to our study, in AST, only 7 patients have been confirmed to have oncogenic PIK3CA variants, of these 3 patients had PIK3CA p.H1047R variant, 3 PIK3CA p.E542K variant, and 1 PIK3CA p.E545K variant (11). Our study is the first to demonstrate PIK3CA mutations also in ECs isolated from AST. Oncogenic PIK3CA variants were detected in our study in the majority of AST lesions (75%, 15/20 patients), supporting thus the finding of Boccara et al. (11) and the importance of oncogenic PIK3CA mutation in AST lesion formation. Additionally, we detected in this study a novel somatic mutation in PIK3CA, p.H1047L, in AST.

Besides genotypically abnormal ECs, other cell types of venous lesions could affect angiogenic phenotype of ECs e.g. via secretion of paracrine growth factors and thus, contributing to lesion formation. We demonstrate here for the first time that TGFA, a known pro-angiogenic growth factor (26,28,29), is upregulated in VM and AST lesions, and in the presence of an oncogenic PIK3CA variant. TGFA and its receptor EGFR located in both intervascular stromal cells and the endothelium. We further demonstrated that patient SCs were able to: i) secrete TGFA and VEGF-A; and ii) transform genotypically normal ECs toward a pro-angiogenic phenotype. Accordingly, our experiments in a modified mouse xenograft model showed an increase in lesion vascularization when genotypically normal fibroblasts were used together with human ECs expressing PIK3CA isoforms. We also demonstrated that afatinib, an irreversible inhibitor of EGFR/ErbB1, ErbB2 and ErbB4, was able to decrease PIK3CA^H1047R^-induced lesion growth and vascularization. As ErbB signaling has been shown to induce activation of RAS/MAPK and PI3K/Akt pathways that are involved in cell proliferation and inhibition of apoptosis (30), targeting of both ECs and intervascular SCs by pharmacological agents could be beneficial to increase treatment response in patients with VM or AST having a fibrous component.

Previously, hypoxic avascular stromal cells were suggested to regulate angiogenesis. For example, cancer-associated fibroblasts can induce tumor initiation, progression and angiogenesis by producing growth factors, proteases, chemokines, and extracellular matrix (39,40). Fibroblasts have also been shown to modulate EC/pericyte migration, and to be crucial for lumen formation. They are also the main source of VEGF-A production in cancer (41). Perturbation of VEGFR signaling is linked to most vascular anomalies and has been demonstrated for example in infantile hemangioma and arteriovenous malformation (42,43). To our knowledge, no study has reported the role of VEGF-A or TGFA in VM or AST. A comparison of various vascular anomalies is needed to understand the possible diagnostic significance of TGFA/EGFR expression in VM and AST.

Current treatment strategies targeting cancer associated fibroblasts aim to: i) inhibit secretion of pro-angiogenic growth factors; ii) reduce accumulation of cells to tumour microenvironment via anti-fibrotic agents; and iii) inhibit expression of lysyl oxidase-like proteins that regulate ECM integrity (44). Whereas cancer cells are considered genetically instable by accruing mutations that allow escape from cellular regulatory mechanisms and enable development of drug resistance, cancer-associated stromal cells are not typically mutated in cancer. In VM, mutations in PIK3CA or TEK genes have been shown to occur solely in EC fraction (36). We also detected PIK3CA mutations only in EC fraction of AST lesions. Besides the clear role of these mutations in ECs driving the lesion formation, we here demonstrate lesion-derived intervascular SCs to be able to secrete pro-angiogenic growth factors that can change genotypically normal EC function and enable angiogenesis. Besides genetic factors, hypoxic environment has been shown to induce overexpression of EGFR in cancer (45) and to upregulate both TGFA and VEGF-A in cancer and ECs (31,32). In addition, depletion of both HIF1A and HIF2A have previously been shown to lead to downregulation TGFA expression (34).

In our study, HIFs and their transcriptional target, VEGF-A, were found to have a strong or moderate positive correlation with TGFA mRNA in patient lesions. TGFA expression was also shown to be upregulated in the presence of oncogenic PIK3CA variant, and common transcriptional targets for patient ECs and HIFs or PIK3CA expressing ECs were detected. As some of the patients used in this study had received sclerotherapy, the treatment may have caused hypoxic environment of cells. To conclude, we have identified, for the first time, involvement of TGFA in vascular lesions and demonstrated the role of fibroblasts in mediating lesion growth and angiogenesis. Targeting of intervascular SCs together with ECs could be beneficial for the treatment of VM and AST with fibrous connective tissue and needs further assessment.

## ACKNOWLEDGEMENT

This study was supported by grants from the Academy of Finland (328835 and 321535 JPL; 287478 and 294073 MUK), Ella and Georg Ehnrooth foundation (JPL), CoE of Cardiovascular and Metabolic Disease (307402, SYH), the ERC grants (GA670951 SYH and 802825 MUK), Sigrid Jusélius Foundation (MUK, SYH), Finnish Foundation for Cardiovascular Research (MUK, SYH, JPL), Jane and Aatos Erkko Foundation (MUK) and Department of Musculosceletal and Plastic Surgery, Helsinki University Hospital (PV). Authors thank Gordon Mills & Kenneth Scott for providing pHAGE-PIK3CA and pHAGE-PIK3CA-H1047R plasmids; National Virus Vector Laboratory (University of Eastern Finland, A.I. Virtanen Institute, Kuopio, Finland) for producing the lentiviral vectors; Single Cell Genomics Core (University of Eastern Finland, A.I. Virtanen Institute, Kuopio, Finland) for preparing and sequencing RNAseq libraries, UEF Cell and Tissue Imaging Unit (University of Eastern Finland, Biocenter Kuopio and Biocenter Finland, Kuopio, Finland) for the support on Confocal imaging and experiments with Incucyte; and the personnel at the Kuopio University Hospital maternity ward (Kuopio, Finland) for providing umbilical cords for HUVEC extraction

## AVAILABILITY OF DATA AND MATERIAL

RNA-seq data has been submitted to NCBI Gene Expression Omnibus under accession number GSE130807 and GSE196311.

## AUTHORS’ CONTRIBUTIONS

HI, SJ and JPL performed research and wrote the manuscript. PV applied for a research permission from Helsinki University Hospital, contacted and informed the patients. PV and ET performed clinical diagnosis, surgery and collected tissue samples for the study. JL did pathological analysis. SJ, HI, SK, MK, JJL and HR performed bead assays or histology. MUK and TÖ did RNA-sequencing and analysis. EA provided control samples. HP and SL did animal experiments. NLK was involved in lentivirus vector work. Vascular Anomaly Team of Helsinki University Hospital (PS, PV, JL, KL, JA) assessed the patients and edited the manuscript. JA provided MRI data. SYH provided materials and reagents for the study. JPL designed and provided materials and reagents for the study.

## SUPPLEMENTARY FILES

S**upplementary Material 1**. NGS experiments

**Supplementary Material 2**. gBlock gene fragments

**Source file for Figure 2 – figure supplement 1**. Raw data for western blot images

**Source file for Figure 6**. Raw data for western blot images

## MATERIALS AND METHODS

### Patient cohort

The multidisciplinary vascular anomaly team of Helsinki University Hospital (HUS) evaluated the patients clinically and radiologically and selected the treatment line. Patient samples were collected in elective surgery in the Department of Plastic Surgery, Helsinki, HUS, Helsinki, Finland. A decision for surgical treatment was based on clinical practices. Patient sample collection was approved by the Ethical Committee of the HUS (Decision No 127/13/03/02/2010 and No 1394/2020). Informed consent was obtained from all patients included in the study. Samples were studied by a pathologist (JL) specialized in vascular anomalies and classified according to ISSVA guidelines by using hematoxylin-eosin staining and immunohistochemistry (Glut-1, CD31, CD34 and D2-40). For DNA, RNA and protein work, tissue samples were taken immediately after resection from the middle of the lesion, snap-frozen in liquid nitrogen and stored at −70⁰C. Optionally, tissue samples were fixed with 4 % paraformaldehyde for immunohistochemical stainings or collected to Dulbecco’s Modified Eagles’ Medium (DMEM; Sigma-Aldrich, St. Louis, MOK, USA) supplemented with 20 % Fetal Bovine Serum (FBS), 20 mM HEPES and antibiotics for cell isolation. Control tissue samples were normal vascular specimens (from mammary artery, n=4; or saphenous vein n=2) from atherosclerotic patients undergoing bypass surgery. After removal, tissue material not needed for a bypass graft was snap-frozen in liquid nitrogen and used for research purposes by approval from the Research Ethics Committee of the Northern Savo Hospital District (Decision No 139/2015).

### Cell culture

Resected patient tissue samples were treated with collagenase type II (Worthington, Lakewood, NJ, USA) for an hour at 37⁰C under agitation. Selection of ECs was performed with CD31 MicroBead Kit and a magnetic column (Miltenyi Biotec, Bergisch Gladbach, Germany) as previously described (46). Patient-derived ECs were cultured on fibronectin/gelatin coated cell culture flasks in Endothelial Cell Growth medium (EGM; Cambrex Biosciences, East Rutherford, NJ, USA) supplemented with 20 % FBS. Patient-derived intervascular SCs from flow-through fraction were maintained in DMEM supplemented with 10 % FBS and antibiotics. HUVECs were isolated from human umbilical cords with approval from the Research Ethics Committee of the Northern Savo Hospital District, Kuopio, Finland (Decision No 341/2015) as previously described (47) and maintained in EGM. Human saphenous vein endothelial cells (HsaVEC) and control human pulmonary fibroblasts (HPF-c) were obtained from PromoCell (3 donors/each, Heidelberg, Germany) and maintained according to manufacturer’s instructions in EGM supplemented with 20 % FBS or in DMEM supplemented with 10 % FBS and antibiotics, respectively. Selection of control ECs was performed by CD31 MicroBead Kit. Patient SCs were characterized prior to experiments by western blot showing to be negative for EC marker CD31, and positive for fibroblast and smooth muscle cell marker vimentin. Additionally, alpha-smooth muscle cell actin, a marker of both myofibroblasts and smooth muscle cells, was detected in 2 cell lines (**Fig. 2 – figure supplement 1D-F**).

### Lentivirus vectors

pHAGE-PIK3CA encoding PIK3CA wild-type (wt; Addgene plasmid #116771; http://n2t.net/addgene:116771; RRID:Addgene_116771) and pHAGE-PIK3CA-H1047R encoding PIK3CA with oncogenic point mutation on p.H1047R (Addgene plasmid #116500; http://n2t.net/addgene:116500;RRID:Addgene_116500) were received as gifts from Gordon Mills & Kenneth Scott (48). Third generation lentiviruses were produced in National Virus Vector Laboratories (NVVL, UEF, Kuopio, Finland). For experiments, HUVECs were seeded onto 6-well plates at a density of 125.000 cells/well and allowed to adhere for 4 hours. Cells were transduced in fresh media with lentivirus vectors expressing PIK3CA WT or PIK3CA p.H1047R with multiplicity of infection (MOI) of 7.5-10. After culturing the cells for 16 h, cells were washed with PBS (Thermo Fisher Scientific, Waltham, MA, USA) and fresh growth medium was added. After 72 h cells were passaged onto new 6-well plates and culturing continued for an additional 72 h after which cells were harvested in Buffer RLT (Qiagen; for RNA-sequencing, RT-qPCR) or used for cell culture experiments.

### RNA-sequencing and gene ontology analysis

Total RNA of ECs was isolated using RNeasy Mini Kit according to manufacturer’s instructions (Qiagen, Hilden, Germany). For patient-derived cell cultures, preparation of RNA-Seq libraries as well as data analysis for differentially expressed genes was performed as previously described (49). Briefly, Poly(A)-RNA was enriched with MicroPoly(A) Purist Kit, fragmented using RNA Fragmentation Reagents (Thermo Fisher Scientific) and purified by running through P-30 column (Bio-Rad Laboratories, Hercules, CA, USA). The 3′ end of the fragmented RNA was dephosphorylated with T4 polynucleotide kinase (PNK, New England Biolabs, Ipswich, MA, USA) followed by heat-inactivation. Dephosphorylation reactions were purified using anti-BrdU beads (SantaCruz Biotech, Heidelberg, Germany) and precipitated overnight. Poly(A)-tailing and cDNA synthesis was performed the next day. After cDNA synthesis, Exonuclease I (New England Biolabs) was used to catalyze the removal of excess oligos. The DNA-RNA hybrid was purified using ChIP DNA Clean & Concentrator Kit (Zymo Research Corporation, Irvine, CA, USA), RNaseH treated and circularized. The libraries were amplified for 11-14 cycles with the oNTI201-primer and a barcode specific primer oNTI200-index. The final product was run on Novex 10% TBE gel, purified and cleaned up as above. The libraries were sequenced on the Illumina Genome Analyzer 2 or HiSeq 2000 according to the manufacturer’s instructions (GeneCore, EMBL, Heidelberg, Germany). RNA-seq was mapped using TopHat (v2.0.7). Poor quality reads were filtered out (minimum 97% of bp over quality cutoff 10) and tag per base value was set to 3. Differentially expressed genes were identified using edgeR (50).

For lentivirus experiments, RNA-Seq libraries were prepared from total RNA using the QuantSeq 3′ mRNA-Seq Library Prep Kit FWD for Illumina (Lexogen, Vienna, Austria) according to the manufacturer’s instructions. The libraries were sequenced with a read length of 68 bp (single end) on an Illumina NextSeq 500 sequencer. The RNA-Seq reads were processed using the nf-core RNA-Seq pipeline (version 3.0) (51) with the GRCh37 genome and the default quantification workflow (STAR aligner for read mapping and Salmon for gene quantification), followed by DESeq2 (version 1.22.2) (52) differential expression analysis.

Each sequencing experiment was normalized to a total of 10^7^ uniquely mapped tags and visualized by preparing custom tracks for the UCSC Genome browser. Clustering results were generated by Cluster 3.0 (53) by normalizing and centering the gene expression tags to range from −1 to 1. The following thresholds were used: FDR < 0.1 (Patient^CD31+^EC vs HUVEC), p-value < 0.05 (Patient ^CD31+^EC vs HsaVEC), FDR-adjusted p-value < 0.1 (PIK3CA^H1047R^ vs PIK3CA^wt^-transduced ECs and PIK3CAmut+ Patient ^CD31+^EC vs Ctrl ECs), RPKM > 0.5 and log2 fold changes > 1.0 and < −1.0. For gene ontology analysis, HOMER 4.3. or the EnrichR web server was used (54–57). Gene Set Enrichment Analysis (GSEA; https://www.biorxiv.org/content/10.1101/060012v3) with a custom gene set calling was used to compare similarity of gene expression pattern between separate experiments. Motif enrichment was analyzed from the merged list of H3K4me2- and H3K27ac-defined enhancers that were located within 100 kb of the transcriptional start site (TSSs) of the differentially expressed genes. The ‘findMotifsGenome.pl’ command in the HOMER software was used with default settings, peak size of 200 bp and motif length of 8, 10 and 12 bases. A random set of genomic positions matched for GC% content was used as background. Enhancer elements enriched for H3K4me2 and H3K27ac marks in HUVECs (data from GSE29611) were generated using the ‘findPeaks’ command in the HOMER software (55) with default settings for ‘style histone’ option: identification of 500 bp regions, 4-fold enrichment over input tag count 4, 0-fold enrichment over local tag count and 0.001 FDR significance. To select the coordinates of enhancers within 100 kb of the TSS ‘mergePeaks’ command with -cobound 1 and -d 100000 were used.

RNA-seq data has been submitted to NCBI Gene Expression Omnibus under accession numbers GSE130807 and GSE196311 (GEO reviewer access tokens; wbivkayaxhojdqp and mbehiikgvtmfryh, respectively). A summary of the NGS samples and gene lists are found in **Table 3** and **Supplementary Material 1**.

### qRT-PCR

Total RNA was isolated from control/patient cells and tissue samples either with RNeasy Mini Kit (Qiagen, Hilden, Germany) or Tri Reagent according to manufacturer’s instructions (Sigma-Aldrich). cDNA synthesis and qRT-PCR were performed using target gene specific Taqman assays (ThermoFisher Scientific, **Table 4**). Amplification of beta-2 microglobulin (B2M; for tissue samples) or glyceraldehyde-3-phosphate dehydrogenase (GAPDH; for ECs, HPF-c and patient SCs) was used as an endogenous control to standardize the amount of RNA in each sample. Detection was performed with StepOnePlus Real-Time PCR System (Applied Biosystems, Foster City, CA, USA).

**Table 4.** Taqman assays used in qRT-PCR analysis.

| Gene | Description | Assay ID |
| --- | --- | --- |
| GAPDH | Glyceraldehyde-3-phosphate dehydrogenase | 4352934E |
| B2M | Beta-2-microglobulin | Hs00187842_m1 |
| TGFA | Transforming growth factor A [assay 1] | Hs00608187_m1 |
| TGFA | Transforming growth factor A [assay 2] | Hs00177401_m1 |
| TGFA | Transforming growth factor A [assay 3] | HsaCEP0053322 |
| ERBB1 | Protein tyrosine kinase ERBB1,<br>epidermal growth factor receptor | Hs01076090_m1 |
| AREG | Amphiregulin | Hs00950669_m1 |
| NRG1 | Neuregulin 1 | Hs00247620_m1 |
| EPGN | Epithelial mitogen, epigen | Hs02385424_m1 |
| VEGF-A | Vascular endothelial growth factor A | Hs00900055_m1 |
| PIK3CA | Phosphatidylinositol-4,5-Bisphosphate 3-Kinase Catalytic Subunit-α | HsaCEP0050716 |

### Recombinant proteins and inhibitors

Cells were seeded on 6-well plates at the density of 200.000 cells/well. When cells reached 80 % confluency they were washed with PBS and synchronized with basal media containing 0.5 % FBS. After 16 h 50 ng/ml of recombinant human (rh)TGFA (Sigma-Aldrich) and/or Afatinib (5 µM, MedChem Express, Monmouth Junction, NJ) was added to the wells. Corresponding concentration of DMSO was used as a control for Afatinib.

### ELISA

Expression levels of TGFA and VEGF-A in cell culture supernatants were measured using Human Quantikine ELISAs (R&D Systems, Minneapolis, MN, USA) according to manufacturer’s instructions. Due to limited availability of patient-derived cells, mRNA expression and protein secretion analysis with patient samples were done on the same wells. Thus, protein concentration measured from cell culture medium was normalized to total RNA extracted from the same well at the same time point.

### Fibrin bead assay

Fibrin bead assay for HUVECs and HPF-c cells has been previously described (35,58). Here, cytodex microcarrier beads were coated with HUVECs and embedded into a fibrin gel. HPF-c cells or, for the first time in this study, patient-derived intervascular stromal^CD31-, vimentin+^ cells were layered on top of the gel with or without rhTGFA, rhVEGF-A (R&D Systems) or their combination (50 ng/ml each). Culturing was continued by changing a fresh EGM ± growth factors every other day. Afatinib (5 µM) or DMSO was added to the wells on day 3 and day 5. On day 7, HPF-c layer on top of the fibrin gel was removed by trypsinization. ECs inside of the gel were fixed, permeabilized and stained with phalloidin-A635 (F-actin, Thermo Fisher Scientific) and DAPI. Imaging was performed using LSM800 (Zeiss). 405/555nm diode lasers were used together with the appropriate emission filters (10x/0.3 PlanApo objective, 512×512 frame size). Image processing and quantitative analysis was performed from 3D-images by ImageJ (59), in a blinded manner by two independent observers. Sprouts containing >1 nuclei were included in the analysis. Segmented vascular area was additionally detected.

### siRNA transfection, followed by imaging of cell growth with IncuCyte

HUVECs expressing PIK3CA p.H1047R or PIK3CA wt were transfected with a Silencer Select siRNA targeting to TGFA (ID: s14053) or negative control siRNAs (#1 and #2 mixed in ratio 1:1; all siRNA oligonucleotides from Thermo Fisher Scientific) as previously described (58). 48h post-transfection, HUVECs were trypsinized, suspended in endothelial basal medium supplemented with 1% FBS, mixed with genetically normal HPF-c (HUVEC-to-HPF-c ratio 8:1) and seeded on 24-well plates at a total density of 15 000 cells/cm^2^ (i.e. 25 000 HUVECs and 3750 HPF-c/well). Cellular growth in the presence of no additional growth factors was monitored using the IncuCyte S3 Live-cell Imaging System (Essen BioSciences Ltd., Hertfordshire, UK). Images were acquired in 3-h intervals, 4 images/well, for a 48-h period using a 10x objective. Mean confluency of the cells at each time point was analyzed, followed by quantitating relative growth rate in each condition based on a slope of the growth curve (**Fig. 4 – figure supplement 2**).

### Immunohistology and whole immunomount stainings

Avidin-biotin-HRP system (Vector Laboratories, Burlingame, CA, USA) with 3’-5’-diaminobenzidine (DAP; Zymed, S. San Francisco, CA, USA) color substrate was used for immunohistochemistry on 4-5 µm thick 4 % PFA-fixed paraffin-embedded sections. Hematoxylin (Vector Laboratories) was used as a background color. Frozen tissue sections (20-30 μm thick) were fixed for double immunofluorescence staining and blocked with a mixture of 1 % BSA and 10 % normal goat serum or with 3 % normal goat serum. Sections were incubated with primary antibodies and Alexa Fluor 488 or Alexa Fluor 594-conjugated secondary antibodies (A11020, A11037; Thermo Fisher Scientific, dilution 1:200). Mounting was performed with Vectashield medium with DAPI (Vector Laboratories). Sections without primary antibodies were used as negative controls. Primary antibodies for all stainings were as follows: rabbit anti-TGFA (HPA042297, Sigma-Aldrich, dilution 1:50 and 1:100), rabbit polyclonal anti-pEGFR clone Tyr845 (07-820, Merck, Kenilworth, NJ, USA, dilution 1:50), rabbit polyclonal anti-EGFR ab (HPA018530, Sigma-Aldrich, dilution 1:100), mouse monoclonal CD31 anti-human clone JC70A (M0823, Agilent Dako, Santa Clara, CA, USA, dilution 1:20 or 1:100) and rabbit polyclonal anti-CD31 ab (NB100-2284, Novus Biologicals, Centennial, CO, USA, dilution 1:50). Imaging was performed by Nikon Eclipse Ni-U microscope (10×/0.3 Plan Fluor or 20×/0.5 Plan Fluor objectives; Nikon, Tokyo, Japan) or by Zeiss LSM800 confocal laser scanning microscope using 405/488/561nm diode lasers together with the appropriate emission filters (20×/0.8 Plan Apochromat, 512×512 or 1024×1024 frame size). Maximum intensity projections were generated using the ImageJ program.

### SDS-PAGE electrophoresis and western blot

Cells treated indicated times with rhTGFA (50 ng/ml) were washed with ice-cold PBS, followed by treatment with lysis buffer [50 mM Tris, pH 7.5, 150 mM NaCl, 1 mM EDTA, 1% Triton X-100, 0.5% sodium deoxycholate, 0.1% SDS, 10% glycerol, 1 mM sodium orthovanadate (Sigma-Aldrich), with protease inhibitors (Roche, Basel, Switzerland)]. Equal amounts of total protein (20 µg) from each sample were loaded on the gel and used for analysis on SDS-PAGE electrophoresis and western blot. Primary antibodies used for the immunodetection were phospho-EGFR ab (2234, CST, MA; dilution 1:1000), total EGFR ab (2646, CST, dilution 1:1000), aSMA (M0851, Dako, dilution 1:250), CD31 (M0823, Dako, dilution 1:500) and Vimentin (M0725, Dako, dilution 1:1000). Horse radish peroxidase (HRP)-conjugated secondary antibodies were purchased from Pierce. Antigen-antibody complexes were detected with PIERCE™ ECL Western Blotting Substrate (Thermo Fisher Scientific) and Gel Dox XR+ Gel Documentation System (Bio-Rad Laboratories).

### Mutational analysis

DNA isolation and ddPCR were performed as previously described (60). Briefly, total DNA was isolated from patient-derived ECs with Tri Reagent according to manufacturer’s instructions (Sigma-Aldrich). DNA isolations from tissue samples were done by lysing 50-100 mg sections of frozen tissue in Hard tissue homogenizing CK28 tubes containing 2.8 mm ceramic beads (Bertin Technologies, Montigny-le-Bretonneux, France) with Precellys homogenizer (Bertin Instruments). Lysed tissues were treated with Proteinase K (Thermo Fisher Scientific) o/n at 50⁰C, followed by a DNA extraction with phenol:chloroform:isoamyl alcohol 25:24:1 (Amresco-inc, Solon, OH). Detection of PIK3CA c.3140A>G (p.H1047R), PIK3CA c.3140A>T (p.H1047L) PIK3CA c.1633G>A (p.E545K) and PIK3CA c.1624G>A (p.E542K) point mutations were performed on the QX200 ddPCR system (Bio-Rad Laboratories) by using PrimePCR ddPCR Mutation assays according to manufacturer’s instructions (Bio-Rad Laboratories). Detection of TEK c.2740C>T (p.L914F) mutation was performed by using custom-design Taqman SNP Genotyping assays [Thermo Fisher Scientific; fwd 5’-CTTCCCTCCAGGCTACTT-3’, rev 5’-AATGCTGGGTCCGTCT-3’, reporter 1 (HEX) 5’-CTTGCGAAGGAAGTCCAGAAGGTTTC-3’, and reporter 2 (FAM) 5’-CTTGCGAAAGAAGTCCAGAAGGTTTC-3’]. Synthetic construct gBlocks Gene Fragments (IDT, Coralville, Iowa, USA; **Supplementary Material 2**) with and without a mutation were designed for each assay and used as positive control DNAs in ddPCR. DNA samples with mutation positive event > 10 and fractional abundance > 0.5 % were considered as mutation positive.

### A modified xenograft model for vascular lesion

Animal experiments were approved by National Experimental Animal Board of Finland (Decision No Esavi-2019-004672) and carried out in accordance with guidelines of the Finnish Act on Animal Experimentation. 2.5×10^6^ HUVECs expressing PIK3CA wt or PIK3CA p.H1047R were suspended in growth factor-reduced and phenol red-free Matrigel (Corning, New York, USA) with or without 0.8×10^6^ HPF-c cells and injected s.c. into both flanks of 6-weeks old female Athymic Nude-Foxn1^nu^ mice (n=18; Envigo, Indiana, USA). For comparison lesion growth with or without oncogenic PIK3CA variant, each mouse had one plug with PIK3CA^H1047R^ ECs and one with PIK3CA^wt^ ECs. Prior injections, mice were randomised to groups receiving either ECs or ECs+HPF-cs, or to be treated with or without afatinib. Lesion size was measured twice a week from d4 onwards with a digital caliper. After lesions reached ∼200µm^3^ in size, 25 mg/kg afatinib was given to mice once daily p.o. for 9 days (diluted in 10% DMSO, 40% PEG300, 5% Tween-80 and 45% saline, MedChemExpress LLC, NJ). Lesions were dissected at d18-20, and lesion size was measured from the dissected explants (NIS-Elements AR). Volume was calculated with the formula volume = (length x width^2^)/2, where the length is the longest diameter and width is the shortest diameter of the lesion. Explants were fixed with 4% paraformaldehyde for 4 hrs and embedded in paraffin. Vascularization, amount of erythrocytes and overall inflammation were evaluated from hematoxylin and eosin or CD31 staining by visual inspection in a blinded manner by one observer (H&E) or 2 independent observers (CD31) and scored on a scale of 0-3, 0 being the lowest score and 3 the highest (0 no vascularization, no erythrocytes, no inflammation/ 1 a few vascular channels, a few vascular channels filled with erythrocytes, mild inflammation/ 2 many vascular channels, many vascular channels filled with erythrocytes, moderate inflammation/ 3 a lot of vascular channels, most of the channels filled with erythrocytes, severe inflammation). EGFR expression was scored (0 no staining, 1 low amount, 2 moderate amount, 3 high amount). Exclusion criteria from the analysis were: i) unsuccessful plug formation; and ii) different anatomical location of the plug.

### Statistical analysis

Results are expressed as means ± SEM. Statistical significance was analyzed using Kruskal-Wallis test with two-stage step-up method of Benjamini, Krieger and Yekutieli to control FDR [**Fig. 1E**: TGFA(1), TGFA(2), AREG; **Fig. 2F**: TGFA in ECs; **Fig. 4E**; **Fig. 2 – figure supplement 1C**: EGFR in ECs; **Fig. 3 – figure supplement 1C-D**; data not normally distributed]; Brown-Forsythe and Welch ANOVA with Dunnet T3 post-hoc test (**Fig. 1E**: NRG1, EPGN; **Fig. 4D**; **Fig. 6B**; data normally distributed but unequal variances); One-way ANOVA with Tukey, Bonferroni or Sidac post-hoc test (**Fig. 4F**; **Fig 6D**; data normally distributed with equal variances); Two-tailed Mann-Whitney U test [**Fig. 2A**; **Fig. 2F**: TGFA in patient SCs; **Fig. 4B**; **Fig. 5G-I** (to compare scoring between each 2 groups); **Fig 7D-E, 7G**; **Fig. 2 – figure supplement 1A**; **Fig. 3 – figure supplement 3A**; **Fig. 4 – figure supplement 1A-B**; **Fig. 5 – figure supplement 1B-C** (to compare scoring between each 2 groups); data not normally distributed]; two-tailed unpaired t-test with Welch’s correction (**Fig. 2G**; **Fig. 7C**; **Fig. 2 – figure supplement 1C**: EGFR in patient SCs; **Fig. 3 – figure supplement 3B**; data normally distributed but unequal variances). p < 0.05 was used to define a significant difference between the groups. Correlation between two markers was analyzed using Spearman rho (**Fig. 3C-E**; data not normally distributed), with values > 0.6 showing strong correlation and values 0.4-0.59 moderate correlation.

